# Engineered mitochondrial A-to-G editors with enhanced efficiency and targeting scope

**DOI:** 10.1101/2024.11.19.623682

**Authors:** Liang Chen, Mengjia Hong, Changming Luan, Meng Yuan, Yiming Wang, Xinyuan Guo, Yue Fang, Hao Huang, Xiaohua Dong, Hongyi Gao, Dan Zhang, Xi Chen, Dihao Meng, Molin Huang, Zongyi Yi, Mingyao Liu, Liangcai Gao, Gaojie Song, Xiaoming Zhou, Wensheng Wei, Dali Li

## Abstract

Mitochondrial base editing is a powerful technology, but the current A-to-G mitochondrial DNA (mtDNA) base editors are reluctant to achieve high efficiency, which is critical for mtDNA engineering. Through laboratory evolution, TadA-8e variants were discovered with substantially increased activity and expanded targeting compatibility, especially for previously unfavorite sequence contexts, when applied to both nuclear and mitochondrial ABEs. Further engineering of the mtDNA editors (eTd-mtABEs) dramatically reduced both DNA and RNA off-targeting effects and enhanced strand-selective A-to-G editing with substitution of DddA to DNA nickases. Moreover, the eTd-mtABEs induced up to 145-fold editing frequencies compared to previous mtDNA adenine base editors in rat cells, and installed targeted mutations in all injected rat embryos with up to 74% efficiency in founders, resulting generation of SNHL and Leigh syndrome models with severe defects. It suggests that eTd-mtABEs are promising mtDNA engineering technology for basic research and for translational studies.

**Highlights:**

1. Engineered TadA variants enhance nuclear and mitochondrial ABE activity in all sequence contexts.
2. Additional mutations in eTd-mtABEs further minimize both DNA and RNA off-targeting effects.
3. Engineered TadA variants are compatible to DNA nickase assisted strand-selective mtDNA editing.
4. eTd-mtABEs enable highly efficient generation of rat mtDNA disease models with severe phenotypes.

## Introduction

Mitochondrial DNA (mtDNA) mutations are one of the primary causes of genetic diseases, whose clinical manifestations are either tissue-specific or multisystemic, with a very high estimated prevalence of approximately 1/5000.^1,2^ Among 95 pathogenic mutations in mtDNA genes, including 13 coding genes, 2 rRNA and 22 tRNA genes, 95% (90/95) of them are point mutations.^1–3^ For a long time, genetic manipulation of mammalian mtDNA has been difficult, which has largely hampered understanding the mechanisms and developing therapeutics for mtDNA diseases, in contrast to the rapid advances in genetic disorders caused by nuclear genes.^4,5^ Despite the availability of a proven method for creating transmitochondrial cytoplasmic hybrid cells (cybrids) to generate cells and mouse models that mimic mitochondrial diseases,^6^ the intricate procedures coupled with the scarcity of technologies to induce targeted mtDNA mutations render the development of disease models, particularly in animals, exceedingly challenging.^5,7^

With the advancement of genome editing technology, mtDNA is targetable through customized nuclease. The proof-of-concept studies have shown that a heteroplasmy shift toward wild-type mtDNA is achievable through site-specific nucleases, specifically mitochondrially targeted restriction endonucleases (mitoREs) and programmable nucleases including zinc-finger nucleases (mtZFNs), transcriptional-activator-like effector nucleases (mitoTALENs) and ARCUS (mitoARCUS), in the transmitochondrial cells or mouse models.^5,8–11^ These encouraging studies shed light on the development of therapeutics for mtDNA diseases. However, since the programmable nucleases are unable to induce specific point mutations, which are the major type of pathogenic mtDNA mutations, targeted engineering of mtDNA to model human mitochondrial disorders remains substantially difficult due to the lack of efficient genetic tools.

Inspired by CRISPR based nuclear base editing strategies, mitochondrial base editors (mitoBEs) have been recently developed by fusing a mitochondrial-localized programmable DNA-binding protein module with a nucleotide deaminase moiety to induce targeted base conversions.^2,12^ With the identification of double-stranded DNA (dsDNA) cytosine deaminase DddA_tox_, DddA-derived cytosine base editors (DdCBEs) and zinc finger deaminases were generated through fusion with uracil glycosylase inhibitor (UGI) and DNA recognition moiety TALEs or ZFNs.^13–20^ By substituting UGI with evolved adenine deaminase TadA-8e variant, mitochondrial adenine base editors (miABEs), named transcription activator-like effector-linked deaminases (TALEDs), were generated to introduce A•T-to-G•C transitions in both strands of dsDNA.^3^ Given that A•T-to-G•C edits theoretically model ∼45% and correct ∼41% of mtDNA diseases,^3,21^ respectively, miABEs hold immense promise for interrogating the function of mtDNA variants and potentially for mtDNA disease therapeutics. To increase the mtDNA editing precision, we and other group have shown that using DNA nickases instead of DddA enhanced the strand-preference base conversions, but the A•T-to-G•C editing efficiency still has room for improvement.^22,23^ A very recent study has developed improved TALED variants with greatly reduced RNA off-target effects and demonstrated generation of a mtDNA disease mouse model, but the phenotype is mild probably due to very limited editing efficiency,^24^ since the clinical manifestation of mtDNA diseases often requires heteroplasmy of mtDNA mutation to exceed a high threshold (typically >50%).^5,25^ As high heteroplasmic mtDNA mutation rate is prerequisite to mimic mitochondrial disorders, highly efficient mitoBEs are urgently demanded in the field.

In this study, through extensive engineering of TadA-8e, we have obtained evolved variants with enhanced editing activity and the targeting scope, especially the RA* (R=A or G) sequence contexts which are unpreferred targets for TadA-8e. Using these super active TadA variants, the engineered TadA-derived mtDNA adenine base editors (eTd-mtABEs) have been developed through fusion with split DddA_tox_ modules or nickases guided by TALE arrays and mitochondrial targeting sequences (MTSs) (Figure 1A). eTd-mtABEs showed enhanced A•T-to-G•C editing frequency comparing to original editors by averaging 6.9-fold (with the TadA-8e-RW variant) or 3.8-fold (with the TadA-8e-RW/V28A variant) in dsDNA- or ssDNA-preferred targeting manner, respectively, with expanded targeting scope and minimized DNA/RNA off-target effects. More importantly, eTd-mtABEs highly efficient installed mtDNA pathogenic point mutations in cells and rats to induce apparent phenotypes for modeling Leber hereditary optic neuropathy (LHON), sensorineural hearing loss (SNHL) and Leigh syndrome. The development of eTd-mtABEs significantly broadens the spectrum of potential applications and enhances the overall versatility of mitochondrial base editing technology.

**Figure 1.**
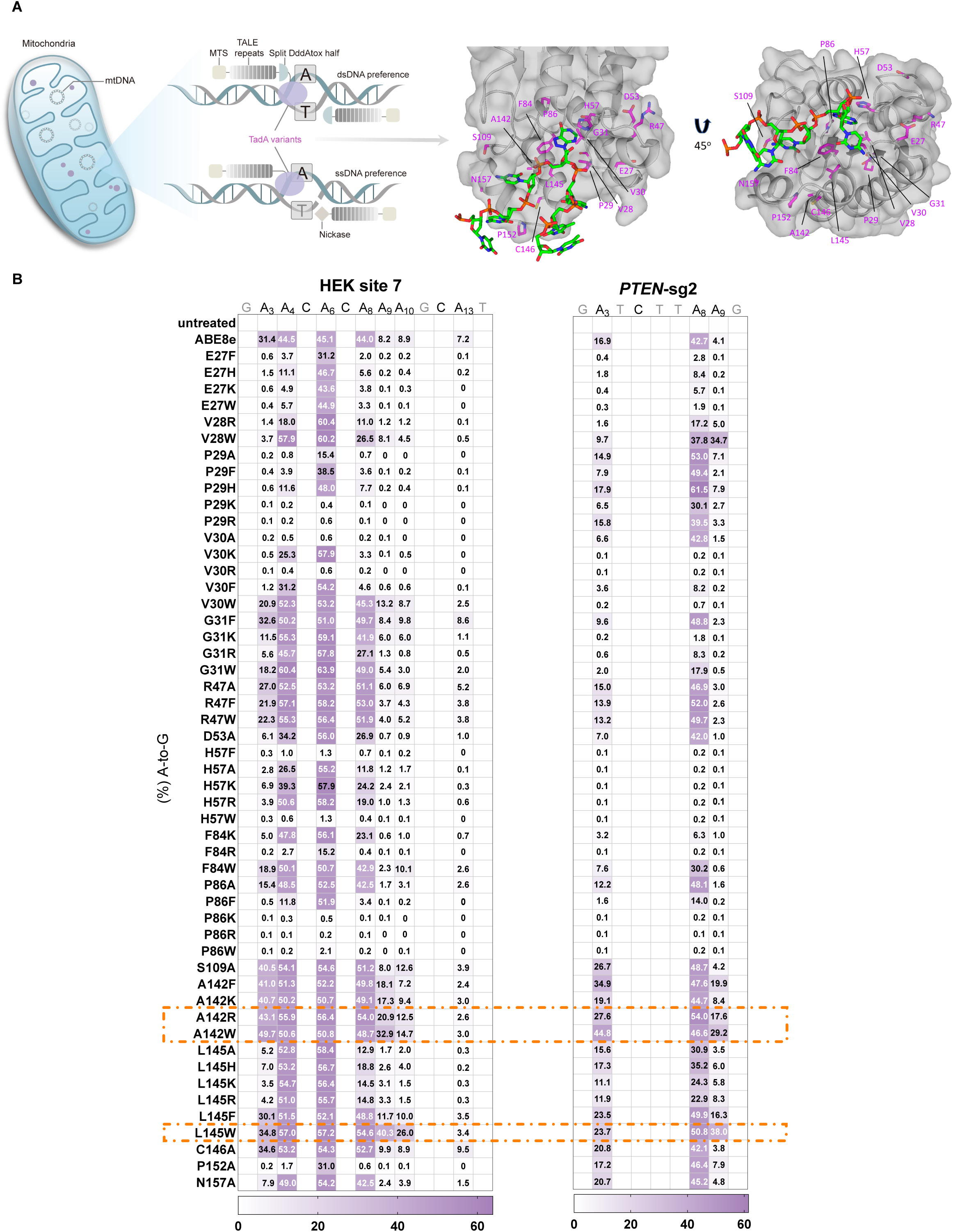
Protein engineering of adenine deaminases. (A) Schematic illustration of eTd-mtABEs using evolved TadA for enhancing dsDNA and ssDNA preferential editing of mitochondria genome. Cryo-electron microscopy structure of TadA-8e in complex with the DNA substrate (right). MTS, mitochondrial targeting sequence; TALE, transcription activator-like effector. (B) Heat maps showing nuclear A·T to G·C editing efficiencies induced by ABE8e and ABE variants at HEK site 7 and *PTEN*-sg2 in HEK293T cells. The A142R, A142W and L145W highlighted in orange dotted boxes are chosen for further evaluation. Data represent the mean of three biologically independent experiments. See also Figure S1.

## Results

### Molecular engineering of adenine deaminases

As the mitochondrial ABEs are less efficient to induce base conversions compared to DdCBEs,^3,13^ we speculate that the activity of adenine deaminase is the most critical index for editing efficiency. Inspired by our previous studies and those of others, which showed that the features of activity, specificity, and substrate selectivity of TadA deaminase can be substantially amended,^26–35^ we attempted to further increase its activity through laboratory evolution to ultimately enhance the editing efficiency of miABEs. To evolve a highly efficient TadA variant, the nuclear editor ABE8e was selected as the starting point due to its ease of construction compared to mitoBEs. Based on the structure of ABE8e in complex with its substrates^36^ (Figure 1A), the amino acids of TadA-8e that are involved in substrate recognition or located in or adjacent to the active pocket were individually substituted, and the editing efficiency was evaluated at an endogenous target (HEK site 7) with scattered adenosines in variant sequence contexts in HEK293T cells (Figure 1B).

Firstly, we examined 51 variants constructed by substituting 16 residues mutated to acquire small side chains, aromatic side chains, polarity with positive charges, or hydrophobic properties (Figure 1B). We found that some variants (for example, E27W, V28R, P29F, P86F, P152A) displayed an extremely condensed editing window (A_6_), and several variants (for example, V28W, G31W, R47F, A142R, L145W) induced a comparable or slightly increased A-to-G efficiency compared to ABE8e in a regular A_4_-A_8_ (PAM sequence as positions 21-23) editing window. Notably, although A_3,_ A_9_ and A_10_ are out of the canonical editing window, the A142R (43% at A_3_, 21% at A_9_, 13% at A_10_) and A142W (50% at A_3_, 33% at A_9_, 15% at A_10_) variants induced a higher ratio of adenine conversions than ABE8e (31% at A_3_, 8% at A_9_, 9% at A_10_). In addition, L145W variant also induced a considerable increase in A-to-G events at position A_9_ and A_10_ (40% and 26%) (Figures 1B and S1A). Similar results were also observed at another target site, demonstrating robust editing activity with an average of 1.9- and 6.9-fold improvement on A_3_ and A_9_ for these 3 variants (A142R, A142W or L145W) (Figures 1B, S1B and S1C). We also noticed that at the evaluated two sites, the activity at all four sequence contexts (NA*, N=A/T/C/G) was increased (Figure 1B). Therefore, these three mutations were selected for further investigation.

### Development of enhanced CRISPR based ABEs

Next, five TadA-8e variants derived ABEs (A142R, A142W, L145W, A142R/L145W and A142W/L145W) were further characterized at 29 endogenous sites (Figure 2A). Compared to ABE8e, all the constructs exhibited increased base editing efficiency to some extent. For example, ABE8e-A142W showed up to 40% and 43% A-to-G editing, whereas ABE8e barely edited adenines with frequencies of 2.8% and 7.7% at A_2_ and A_8_ of *EMX1* site 1, respectively. Higher base conversion rates were induced by ABE8e-A142R/L145W (ABE8e-RW) compared to ABE8e (55% versus 21% at A_9_) at another target (*PD-1*-sg13). Overall, ABE8e-RW exhibited the highest average editing activity among these five constructs. It showed a dramatic elevation of editing activity at position A_-1_-A_3_ (averaging 4.9-fold) and A_8_-A_11_ (averaging 2.6-fold) in comparison to ABE8e (Figures 2B, 2C and S2A). An up to 20-fold increase in adenine conversions was observed for ABE8e-RW at A_10_. Even within the conventional editing window (A_4_-A_7_) which is considered nearly saturated in ABE8e,^30^ the activity of ABE8e-RW exhibited an average 1.2-fold increase (Figures 2B and 2C). Previous studies showed that ABEs, including ABE7.10, ABE8s and ABE8e, had a preference for YA* (Y=T/C) motif context, especially outside the conventional editing window.^26,30,31,37^ After carefully analyzing A-to-G conversion fold changes from A_-1_-A_11_ position in the above 29 sites, we found that, compared to ABE8e, ABE8e (especially the A142W and RW) variants showed enhanced activity in all sequence contexts, with more efficient adenine conversions (up to 4.5-fold on average) in RA* (Y=G/A) motifs which were considered inefficient contexts for TadA7/8-derived ABEs (Figures 2D and S2B).

**Figure 2.**
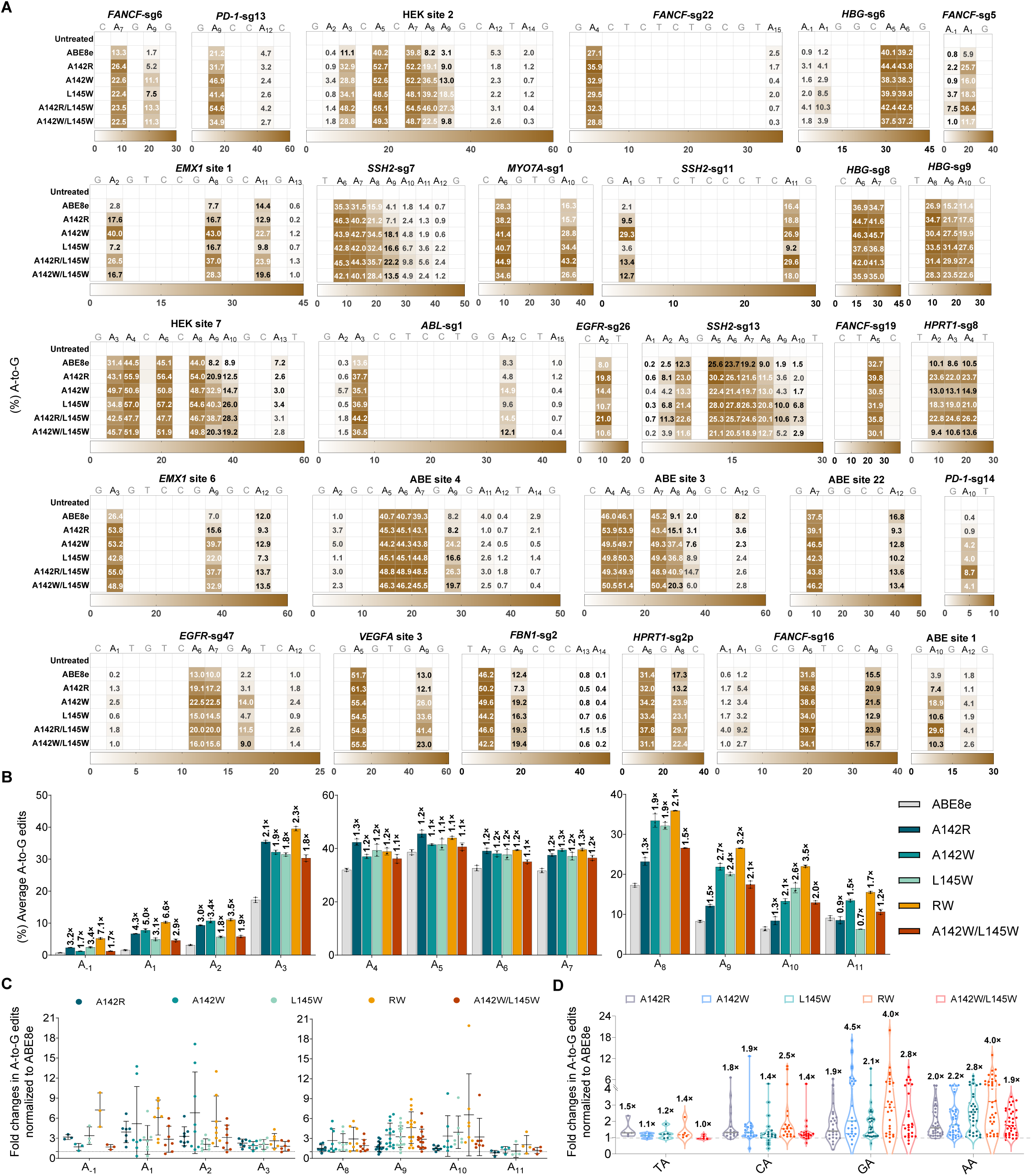
Characterization of enhanced CRISPR derived ABEs. (A) Heatmaps showing nuclear A-to-G editing efficiencies by indicated ABEs at 29 endogenous target sites in HEK293T cells. Data are shown as the mean from n = 3 biologically independent samples. (B) Average A-to-G edits of indicated ABEs in the protospacer at 29 target sites in (A). Fold changes of ABE variants/ABE8e on average A-to-G editing are shown on the top of bars. Data are shown as the mean ± s.d. from n = 3 biologically independent samples. (C) Dot plot represents fold changes in A-to-G edits of indicated ABEs normalized to ABE8e at the positions outside the conventional editing window. Data represent the mean ± s.d., and each dot represents the mean of three biologically independent samples. (D) Violin plots representing fold changes in A-to-G edits of the indicated ABEs normalized to ABE8e in all sequence contexts (TA, n=5; CA, n=20; GA, n=27; AA, n=36) at 29 endogenous genomic loci in (A). Fold changes on average are shown on top of each column. Each dot represents the mean of three biologically independent samples. See also Figure S2.

Previous studies have demonstrated that ABEs also induce cytosine bystander edits.^33,34,38^ Interestingly, except for the A142R variant, the other four evolved ABE8e variants only induced minimal indels and very mild cytosine mutations (less than 1.2% on average) compared to ABE8e (less than 2.9% on average) at the evaluated sites (Figures S2C and S2D). It suggests that L145 residue is critical for distinguishing between adenine and cytosine substrates and reducing cytosine bystander editing, which is consistent with our previous study.^33^ These results indicate that the introduction of substitutions at the 142 and/or 145 residues of TadA-8e considerably enhances editing activity, targeting scope and reduces cytosine bystander editing of nuclear ABEs.

### Robust mtDNA editing with engineered TadA variants

The dramatically increased deaminase activity and targeting scope of TadA-8e variants encouraged us to investigate their potential for mitochondrial A-to-G base editing. The evolved mtDNA adenine base editors were constructed through fusion of wide-type DddA_tox_ halves split at G1397 and TadA-8e adenine deaminase (AD) variants along with MTS and TALE arrays (Figure S3A), resulting in a pair of L-1397C-AD variants/R-1397N or L-1397N/R-1397C-AD variants (eTd1-mtABEs). These eTd1-mtABEs were targeted to three mtDNA sites previously examined with sTALED in HEK293T cells.^3^ HTS data showed that sTALED (AD in L-TALE) induced A-to-G editing frequencies ranging from 6.5-29% at positions 9-18 (the spacer region of two TALE-binding sites) on *ND5* site 2. Excitingly, all of eTd1-mtABEs achieved much higher adenine conversions from 14-71%, and the A142W variant was more active than the others (28-71%) at this site (Figure 3A). Similar results were also obtained from the other construction with AD fused to R-TALE array, and the editing frequency within the same editing window was 52% on average in eTd1-mtABE-A142W treated cells, but the sTALED only induced 24% on average (Figure S3B). When targeted to the *ND1* site 1, sTALED poorly catalyzed A-to-G conversions (7%) when AD was fused to the R-TALE array, while higher efficiency was observed in L-1397C-AD construction (26-39%, averaging 34%). Similarly, eTd1-mtABE-A142R/L145W (eTd1-mtABE-RW) with either in R-TALE (21-35%, averaging 26%) or in L-TALE (54-79%, averaging 66%) construction showed a dramatic increase of adenine edits compared to the counterparts of sTALED (Figures 3B and S3C). In contrast to sTALEDs with averaging 17% editing at *ND4* site 1, all eTd1-mtABEs extended the editing window from A_5_-A_8_ to A_5_-A_10_ with averaging 28-44% editing efficiency regardless of AD locations (Figures 3C and S3D). In addition to tremendously elevated mtDNA on-target editing efficiency, eTd1-mtABEs showed high product purity and rarely induced indels similar to sTALEDs^3^ (Figures S3E-S3J).

**Figure 3.**
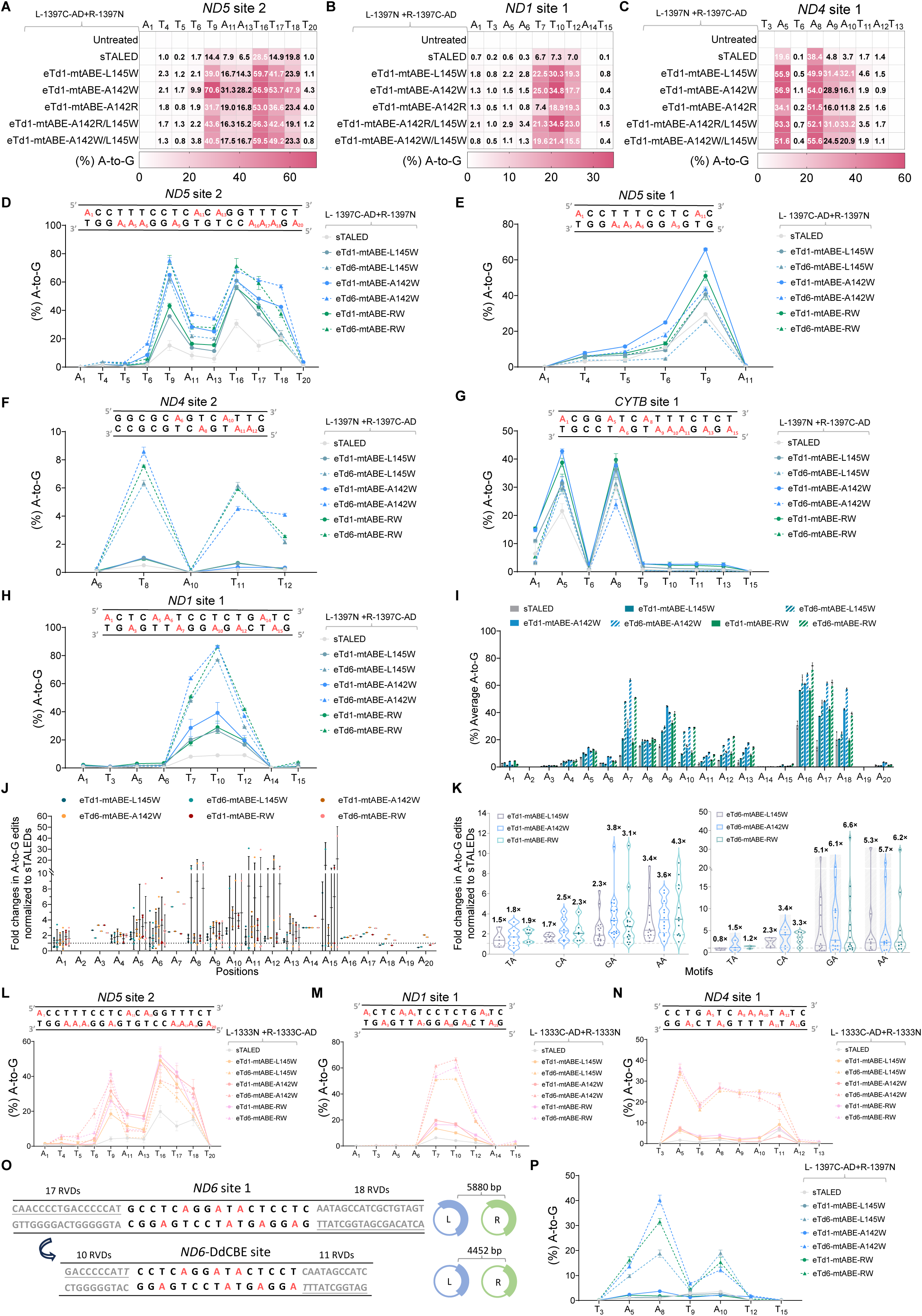
Evolved TadA variants improve A·T-to-G·C editing in mtDNA. (A-C) Heatmaps showing mitochondrial A·T-to-G·C editing frequencies induced by sTALEDs and eTd1-mtABEs at *ND5* site 2 (A), *ND1* site 1 (B) and *ND4* site 1 (C) in HEK293T cells. Data represent the mean of three biologically independent replicates. (D-H) Editing frequencies of indicated editors at *ND5* site 2 (D), *ND5* site 1 (E), *ND4* site 2 (F), *CYTB* site 1 (G) and *ND1* site 1 (H) in HEK293T cells. (I) Average mitochondrial A-to-G editing frequencies induced by indicated eTd-mtABEs. (J) Fold changes in A-to-G edits mediated by indicated eTd-mtABEs normalized to sTALEDs (dashed line). Data represent the mean ± s.d., and each dot represents the mean of three biologically independent replicates. (K) Violin plots representing fold changes in A-to-G edits of indicated eTd1-mtABEs (left; TA, n=5; CA, n=6; GA, n=17; AA, n=12) and eTd6-mtABEs (right; TA, n=3; CA, n=5; GA, n=15; AA, n=10) normalized to sTALEDs (dashed line) in all sequence contexts at target sites in (A-H). Fold changes on average are shown on top of each column. Each dot represents the mean of three biologically independent replicates. (L-N) Mitochondrial A-to-G editing of sTALEDs and eTd-mtABEs with DddA_tox_ split at G1333 at *ND5* site 2 (L), *ND1* site 1 (M) and *ND4* site 1 (N) in HEK293T cells. (O) The comparison of *ND6* site 1 and *ND6*-DdCBE site (left). The length of coding regions for mitochondrial A-to-G editors using corresponding RVDs (right). TALE-binding sites contained a mismatched terminal RVDs are shown in underline. (P) Mitochondrial A-to-G editing frequencies of sTALED and eTd-mtABEs at *ND6*-DdCBE site. For (I) and (J), data were obtained from 5 sites in (D-H). For (D-I), (L-N) and (P), data represent the mean ± s.d. of n=3 biologically independent replicates. See also Figures S3 and S4.

We thought that the recently reported highly active DddA variants^39^ might have better ability to unwind mtDNA which would further improve eTd-mtABE efficiency. eTd-mtABEs with DddA6 variant (eTd6-mtABEs) induced up to 76% A-to-G edits and displayed averaging 1.6- and 4.3-fold improvement than original eTd1-mtABEs and sTALED at *ND5* site 2, respectively (Figure 3D), but introduction of DddA11 variant almost abolished editing activity (Figure S4A). After evaluation of four more targets, we found that the editing frequency of eTd1-mtABEs and eTd6-mtABEs were much higher (up to 11-fold increase and 36-fold increase, respectively) than sTALEDs (Figures 3E-3J). However, eTd6-mtABEs was not always more efficient than eTd1-mtABEs. For example, eTd6-mtABEs showed higher efficiency up to 87% at two targets (*ND1* site 1 and *ND4* site 2), but lower efficiency at *CYTB* site 1 and *ND5* site 1 compared with eTd1-mtABEs (Figures 3E-3H). It suggests that the cytosine deaminase activity of DddA variants is probably not necessarily consistent with the DNA unwinding capability to facilitate TadA deamination. These data suggest that all eTd1-mtABEs exhibited much higher activity compared to sTALEDs regardless of the location of the evolved deaminase, DddA versions and their split positions. Notably, consistent with the results in nuclear ABE8e variants, eTd-mtABEs showed substantially increased activity in all sequence contexts compared to sTALEDs, especially in the RA* contexts where eTd1-mtABEs and eTd6-mtABEs showed up to an average of 4.3-fold and 6.6-fold increase (up to 36-fold at a GA motif), respectively, suggesting highly efficient mtDNA A-to-G editing with an expanded targeting cope (Figure 3K).

### Compatibility of TadA variants to split orientations and short TALE arrays

The split orientation of DddA_tox_ is one of the determinants affecting the editing efficiency of mitoBEs, and higher mutation rates were achieved using G1333-split DddA_tox_ halves for some targets.^3,13^ sTALEDs, eTd1-mtABEs and eTd6-mtABEs with DddA_tox_ split at G1333 were tested at three sites. The average conversion rates of adenines within the spacers of all three targets were 5.4% for the sTALEDs-G1333 construction but it was higher for the sTALEDs-G1397 construction (13%) (Figures 3A-3C and 3L-3N). Compared to the sTALEDs-G1333 (averaging 5.4%), eTd1-mtABEs-G1333 (averaging 15%) and eTd6-mtABEs-G1333 (averaging 32%) catalyzed much higher A-to-G conversions. For example, eTd6-mtABE-A142W and eTd6-mtABE-RW variants showed higher activity (averaging 33% and 34%, respectively) with frequencies up to 67%, even surpassing their counterparts that utilized DddA-G1397 split orientation at some positions (Figures 3A-3C and 3L-3N).

Compared to DdCBEs which usually consist of ∼29 RVD modules, miABEs normally require more RVD modules (∼38 RVDs) to achieve efficient editing.^3,13,39^ As our TadA-8e variants exhibited very high activity, we speculated that super-active deaminase activity would be tolerant of shorter TALE arrays, which would benefit the delivery of the editors. The *ND6* site 1 was used for evaluation, since previous reports demonstrated TALEDs with 35 RVD modules and DdCBEs with 21 RVD modules efficiently generated targeted editing^3,13^ (Figure 3O). When using the construction of conventional 35 RVD modules, sTALEDs induced efficient A-to-G editing, whereas the eTd-mtABEs showed much higher activity (Figure S4B). Once using the construction of 21 RVD modules, sTALEDs almost lost the activity (<4%). In contrast, eTd-mtABEs, especially the eTd6-mtABE-A142W and eTd6-mtABE-RW, induced robust A-to-G conversions with frequencies up to 40% (Figure 3P). Together, these data suggest that TadA variants are very efficient in supporting the fusion of diverse split orientation of DddA_tox_ halves, and are compatible for short TALE arrays which reduces the size by about 25% (∼1400bp) while achieving efficient editing at the tested site, indicating an advantage for delivery.

### Off-target analysis of evolved mitochondrial A-to-G editors

To profile the off-target activity of eTd-mtABEs, we first evaluated whether they would induce nuclear DNA editing at previously identified sites with similar TALE arrays.^3^ Like TALEDs, eTd-mtABEs induced a background level of off-target mutations in nuclear genome (Figure S5A). Next, whole mitochondrial genome sequencing (WGS) of HEK293T cells transfected with sTALEDs, eTd-mtABEs and TALE-free control was performed to test the TALE-independent off-target editing. Compared to sTALEDs, eTd-mtABEs exhibited comparable or slightly increased DNA off-target editing levels without sequence preference (Figures 4A, S5B and S5C). Inspired by studies on nuclear ABEs, which showed that substitutions on critical residues increased substrate selectivity and dramatically reduced off-target edits,^27–29,32,33,40^ we generated 28 constructs in combining eTd6-mtABE-A142W or eTd6-mtABE-RW with other mutations. After evaluation at two endogenous targets, three constructs (RW/V28A, RW/V106W and V82S/A142W/Q154R) showed consistently comparable or increased A-to-G edits (Figure 4B). Then, five representative DNA off-targeting sites identified by the above mitochondrial WGS were used to evaluate the off-target effects of above triple mutant variants in eTd1-mtABE and eTd6-mtABE constructions. HTS data revealed that all six constructs substantially reduced DNA off-target editing comparing to sTALED (Figure 4C).

**Figure 4.**
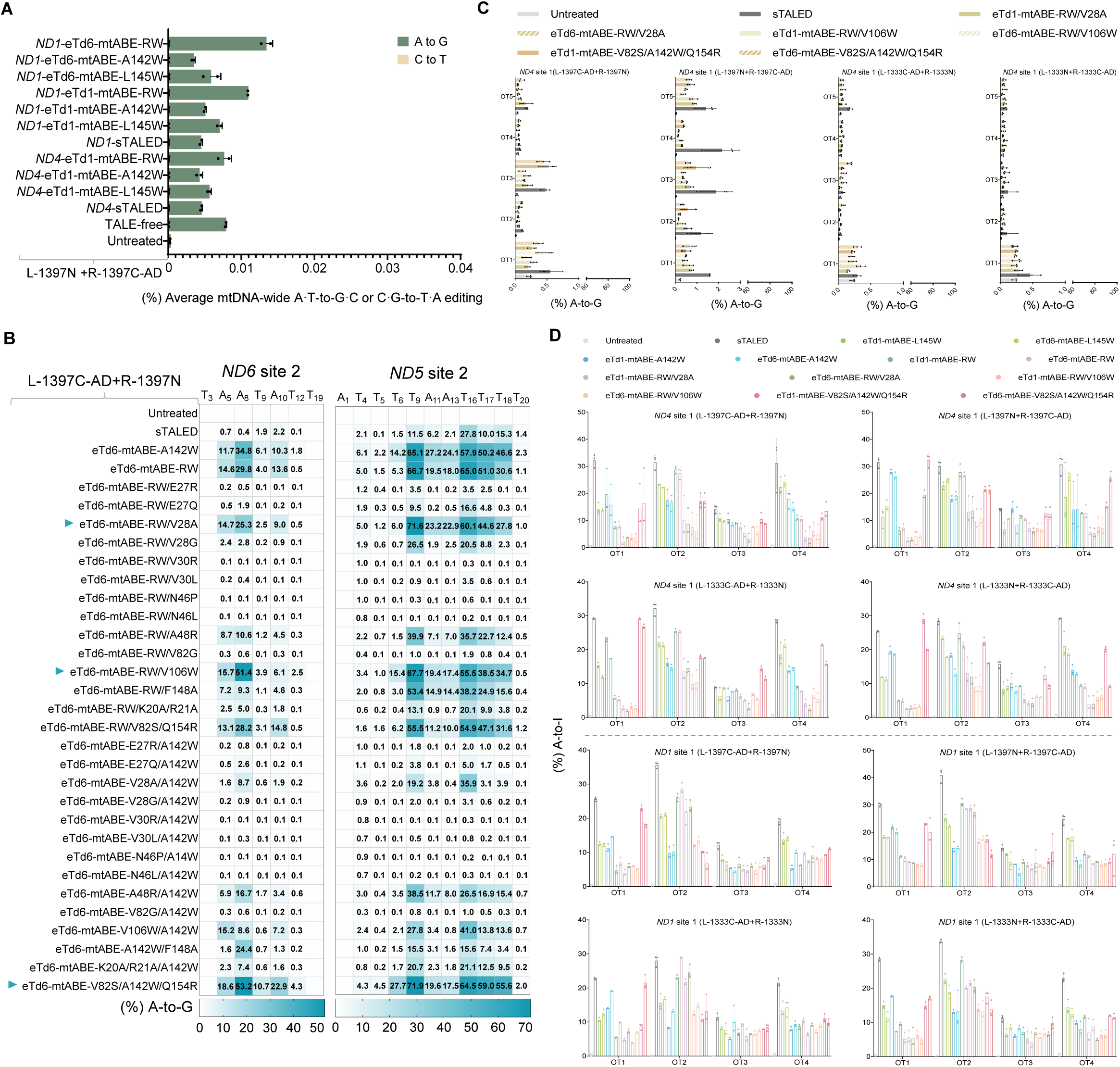
DNA and RNA off-target assessment of eTd-mtABEs. (A) Average percentage of mitochondrial genome-wide off-target editing for sTALEDs and eTd-mtABEs targeting *ND4* and *ND1* sites, respectively. Data represent the mean ± s.d. of n=2 biologically independent replicates. (B) Comparison of A-to-G efficiencies by sTALED and eTd6-mtABE variants at the *ND6* site 2 and *ND5* site 2 in HEK293T cells. eTd6-mtABE variants marked with blue arrows are chosen for further evaluation. Data represent the mean of three biologically independent replicates. (C) DNA off-target editing frequencies induced by sTALEDs and eTd-mtABEs at five representative DNA off-target sites identified by mitochondrial genome-wide sequencing. (D) Bar plots showing RNA off-target editing frequencies induced by sTALEDs and eTd-mtABEs at four high-frequency RNA off-target sites identified by transcriptome-wide sequencing. For (C) and (D), data represent the mean ± s.d. of n=3 independent replicates. See also Figure S5.

A most recent study showed that the original mitochondrial A-to-G editors induced considerable RNA off-target edits.^24^ To evaluate this issue, four previously identified high-frequency RNA off-target sites^24^ were selected for HTS analysis using four different constructs targeting two endogenous sites (*ND4* site 1 and *ND1* site 1). In contrast to TALEDs which showed 9-41% RNA off-target editing, all eTd-mtABEs, especially two variants (RW/V28A and RW/V106W) induced minimized random RNA off-target editing (Figure 4D). The above data suggest that these eTd-mtABE variants (especially RW/V28A and RW/V106W) exhibit increased on-target activity but dramatically reduced off-target events compared to sTALED both at the DNA and RNA levels.

### Enhancing precise mtDNA editing with engineered TadA variants

Recent studies, including ours, demonstrated that fusion of DNA nickases instead of DddA enabled strand-preferred mitochondrial base editing, but their limited activity would hinder their applications.^3,13,22,23,39^ As the deaminase is a critical limiting component of this system, we supposed that highly efficient TadA variants could enhance their performance. Five engineered TadA variants were individually introduced into mitoABE^MutH^, a mitochondrial A-to-G editor with specific nicking preferences (5′-GATC-3′) to achieve selective editing at unnicked mtDNA strand.^22^ Editing results at *ND4* site 1 showed that these variants kept the strand-biased editing feature with obviously increased activity. For example, within the editing window (A_5_-A_10_), the TadA-8e-RW variant showed averaging frequencies of up to 49% which was about 2-fold of original mitoABE^MutH^ (mean 25%), and the efficiency of the highest editing position was also substantially increased (62% vs 39%) (Figure 5A). At the hardly edited position for mitoABE^MutH^ (9.2% at A_9_), the RW variant displayed robust A_9_-to-G edits (39%). Importantly, two precise variants (RW/V28A and RW/V106W) also exhibited very high editing activity comparable to RW variant. Moreover, substantial improvement of efficiency by TadA-8e variants were also applicable to mitoABE^Nt.BspD6I(C)^ which recruited truncated Nt.BspD6I nickase without sequence context constraints^22^ (Figures 5B-5D). Compared to mitoABE^Nt.BspD6I(C)^, TadA-8e variants derived editors achieved much higher A-to-G edits with averaging 3.2-fold improvement at all edited adenines of three tested sites. Even at the highest positions of each site, evolved TadA-8e variants showed over 2-fold increase, such as on the highest position at *RNR2* site 1 (25% versus 7%), *ATP6* site 1 (3.7% versus 1.7%) and *ND1* site 1 (44% versus 22%), and kept a low level of undesired editing at the opposite strand (Figures 5B-5D). These results indicate that the evolved TadA-8e variants considerably enhanced the efficiency of strand-biased mitochondrial A-to-G editing without affecting editing precision.

**Figure 5.**
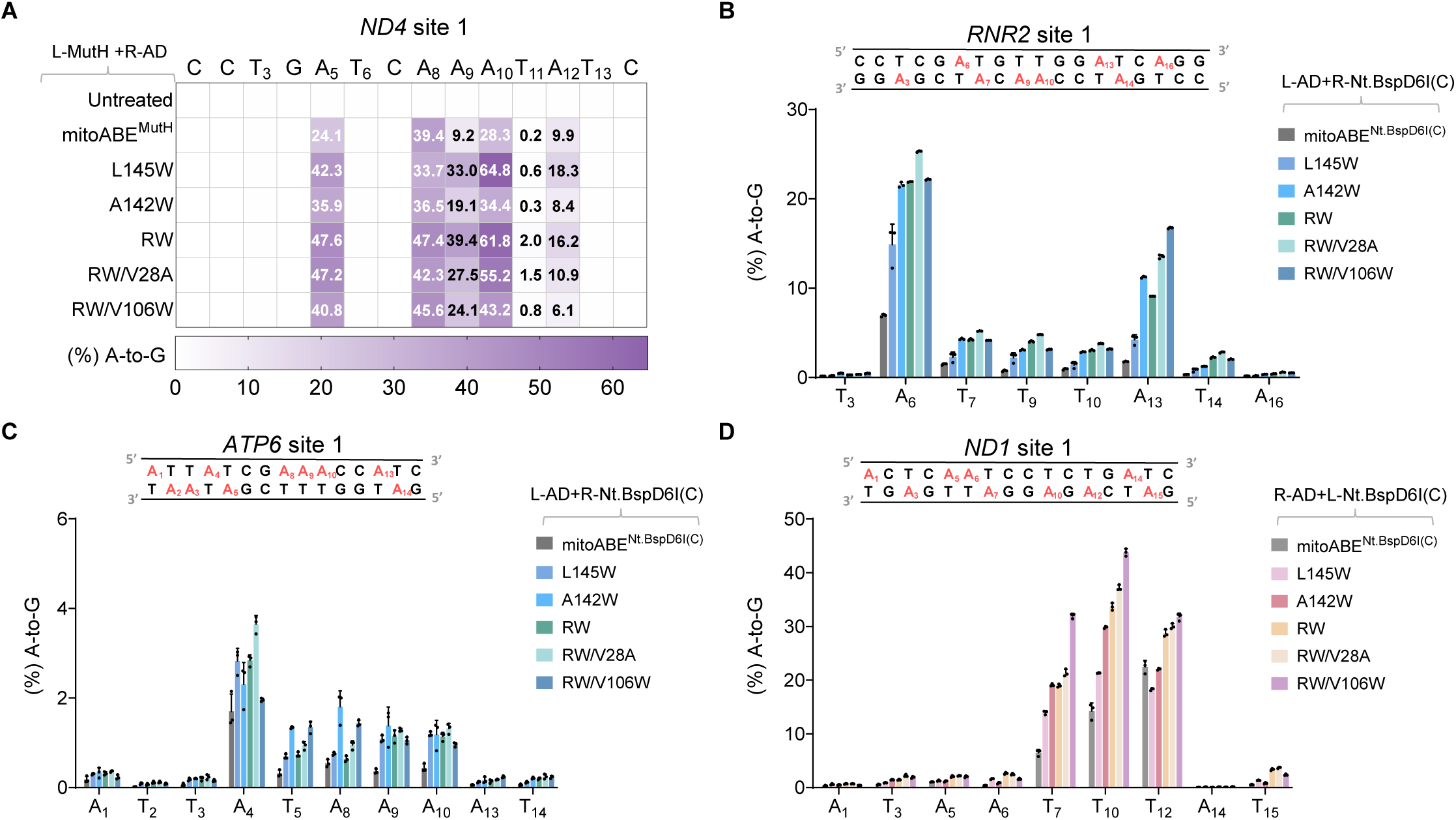
TadA variants enhanced strand-biased mtDNA editing. (A) Heatmap showing A-to-G editing efficiencies of canonical mitoABE^MutH^ and mitoABE^MutH^ constructs with indicated TadA variants at *ND4* site 1 in HEK293T cells. Data represent the mean of three biologically independent replicates. (B-D) A-to-G editing efficiencies of canonical mitoABE^Nt.BspD6I(C)^ and mitoABE^Nt.BspD6I(C)^ constructs with indicated TadA variants at *RNR2* site 1 (B), *ATP6* site 1 (C) and *ND1* site 1 (D) in HEK293T cells. Data represent the mean ± s.d. of n=3 independent replicates.

### Introduction of pathogenic SNVs via eTd-mtABEs in human cells

To test whether eTd-mtABEs induced mtDNA mutations could cause phenotypic effects, we attempted to evaluate the potential of eTd-mtABEs to mimic pathogenic SNVs in human cells. As m.14484T>C and m.14487T>C mutations in *ND6* gene are implicated in LHON and Leigh syndrome,^41–44^ respectively, eTd1-mtABEs and eTd6-mtABEs with G1397 or G1333 orientation were constructed to target the *ND6* site 3 covering the pathogenic mutations (Figure 6A). sTALEDs-G1397 induced less than 15% and 3.8% editing at the desired T_8_ and T_11_ positions which were corresponding to above two pathogenic SNVs, and the G1333-orientation showed even lower activity (less than 6.7% and 2.9%, respectively). For instance, the eTd1-mtABE-RW and the accurate eTd1-mtABE-RW/V28A variants supported highly efficient desired T_8_ and T_11_ editing with either G1397 orientation (41-48% and 23-30%, respectively) or G1333-orientation (26-33% and 11-21%, respectively) (Figure 6B). In addition, compared with sTALED (3%), up to 26% of edited alleles contained simultaneous m.14484 and m.14487 T-to-C mutations when using eTd1-mtABE-RW, indicating its high activity (Figures S6A and S6B). Then, we evaluated the phenotypic consequences of these pathogenic SNVs. Compared to untreated cells and cells containing inactive monomer L-1397N, higher levels of reactive oxygen species and lower ATP levels were observed in cells treated with eTd1-mtABE-RW (Figures 6C and 6D). As the cellular phenotype is highly dependent on mtDNA mutation rates, these data suggest that eTd-mtABEs can achieve highly efficient editing to reach the mutation threshold necessary to induce phenotypes in cells, indicating that they are optimal tools for generating cellular models of mitochondrial diseases.

**Figure 6.**
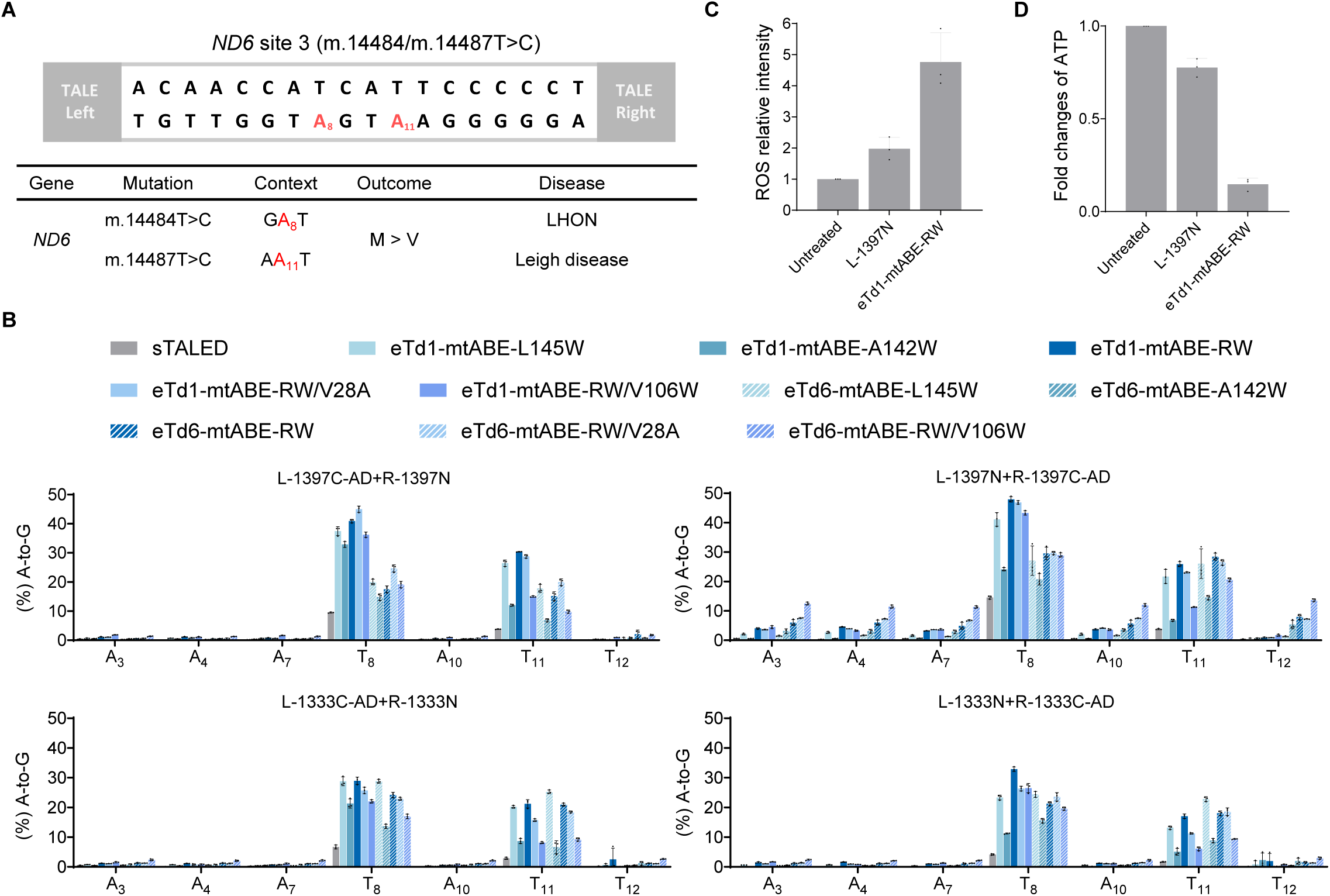
Application of eTd-mtABEs to install pathogenic mutations in human cells. (A) Using eTd-mtABEs to install disease-associated mutations in human mtDNA. Two pathogenic mutations located in target spacer region are shown in red. M, Methionine; V, Valine; LHON, Leber hereditary optic neuropathy. (B) A-to-G editing frequencies of sTALEDs and eTd-mtABEs with DddA_tox_ split at G1397 or G1333 targeting *ND6* site 3. (C) The level of intracellular ROS in HEK293T cells treated with L-1397N, eTd1-mtABE-RW (L-1397N+R-1397C-AD) targeting *ND6* site 3. (D) Fold changes of the intracellular ATP in HEK293T cells treated with L-1397N, eTd1-mtABE-RW (L-1397N+R-1397C-AD) targeting *ND6* site 3 using untreated groups as normalized control. For (B-D), data represent the mean ± s.d. of n=3 independent replicates. See also Figure S6.

### eTd-mtABEs enable mitochondria disease modeling in rats

Animal models of mtDNA diseases are invaluable resources for both basic and translational studies. We attempted to use hyperactive eTd-mtABEs to generate inheritable rat disease models, as the only animal model created via A-to-G conversion associated with mtDNA diseases was generated in mice, albeit with relatively low efficiency.^24^ Three pathogenic mtDNA SNVs, m.7510/7511 T-to-C mutations in *TRNS1* gene and m.14487 T-to-C mutation in *ND6* gene, cause sensorineural hearing loss (SNHL)^45,46^ and Leigh diseases,^43,44^ respectively (Figures 7A, 7B and S7A). Firstly, we constructed five eTd-mtABEs along with sTALEDs to target the pathogenic T_9_ and T_10_ positions (corresponding to rat m.6929/6930 positions) of *TRNS1* site 1 in PC12 cells (Figure 7C). In contrast to sTALEDs which hardly induced desired T_9_ and T_10_ conversions (0.8% and 0.9% respectively), all eTd-mtABE constructs induced substantially higher base transitions. eTd6-mtABE-RW exhibited superior activity representing an up to 145-fold increase compared to sTALED, achieving up to 49% and 42% editing at the T_9_ and T_10_ positions respectively (Figures 7D and S7B). Then, eTd6-mtABE-RW mRNA was injected into rat zygotes, and amazingly, 100% (54/54) of F0 pups bearing desired T_9_-to-C (averaging 27%, 2.7-44%) and T_10_-to-C mutation (averaging 27%, 3.8-44%) were obtained (Figure 7E). Notably, 13 of the founders carried simultaneous pathogenic m.6929 and m.6930 T-to-C mutations with frequency of over 25% (Figures 7F and S7C). Auditory brainstem response (ABR) assays across frequencies at 3, 10, 15, and 20 kHz from 4-week-old F0 founders demonstrated that ABR thresholds of 8 founders were significantly increased at frequencies of 3 kHz and 20 kHz than wild-type rats (Figure 7G), with severer phenotypes in female founders (Figure S7D), suggesting successful generation of mitochondrial disease model of SNHL with hearing disorders.

**Figure 7.**
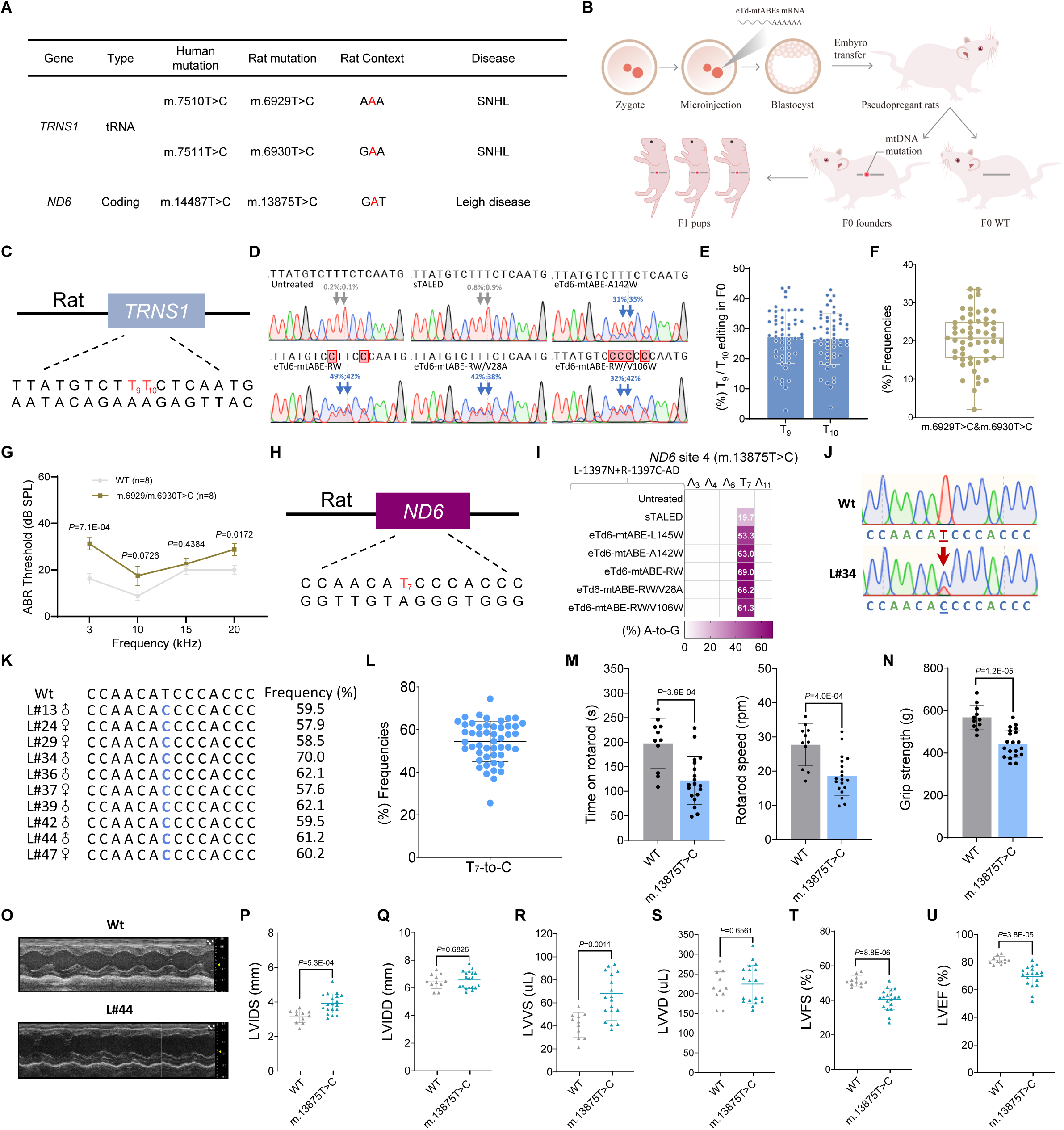
Mitochondrial disease models generated by eTd-mtABEs in rats. (A) Disease-associated mutations in human mtDNA and corresponding rat mtDNA. Pathogenic mutation sites were highlighted in red. SNHL, sensorineural hearing loss. (B) The workflow for eTd-mtABE-mediated generation of disease models in rats carrying pathogenic mutations. (C) The target sequence of the rat mitochondrial *TRNS1* gene is shown. Two pathogenic mutation sites (T_9_ and T_10_) are highlighted in red. (D) Sanger sequencing chromatogram of *TRNS1* site 1 edited by indicated editors (L-1333N+R-1333C-AD) in PC12 cells. Arrows indicate desired editing positions. Data represent the mean of three biologically independent replicates. (E) Desired T_9_-to-C or T_10_-to-C editing frequencies in F0 rats. Each data point represents an individual rat (n = 54). (F) Bar plots represent frequencies of alleles with simultaneous m.6929T>C&m.6930T>C editing. (G) Auditory brainstem response (ABR) thresholds of wild-type rats and representative rats with m.6929/m.6930T>C at 4 weeks old. SPL, sound pressure level. Data represent the mean ± s.d., and *P* values were calculated two-tailed Student’s t-tests. (H) The target sequence of the rat mitochondrial *ND6* gene is shown. The pathogenic position is shown in red. (I) Heatmap showing frequencies of mitochondrial A-to-G conversion treated with indicated editors targeting the *ND6* site 4 in PC12 cells. Data represent the mean of three biologically independent replicates. (J) Sanger sequencing chromatogram of *ND6* site 4 in a wild-type rat and a representative F0 founder. The red arrow indicates the target site. (K) Genotyping of representative F0 founders. The desired T_7_-to-C conversion is highlighted in blue. Frequencies of mutant alleles were determined by HTS. (L) Desired T_7_-to-C editing frequencies of F0 founders with m.13875T>C mutation. Each data point represents an individual rat (n = 49). (M) Motor coordination of wild-type rats and F0 founders at 11 weeks old was evaluated by rotating rod test, with time on rotarods (left) and rotarod speed (right). (N) Grip strength of mutant F0 rats with m.13875T>C was measured by grip strength test compared to wild-type rats at 11 weeks old. (O) Snapshots of M-mode of echocardiographic measurements including a wild-type rat and a representative rat. (P-U) Echocardiographic measurements of wild-type rats and F0 founders at 12 weeks old. Tests include left ventricular diameter at end-systole and end-diastole (LVIDS, LVIDD) (P and Q), left ventricular volume at end-systole and end-diastole (LVVS, LVVD) (R and S), left ventricular percent fractional shortening (LVFS) (T) and left ventricular ejection fraction (LVEF) (U). For (E-F), (L), (M-N) and (P-U), data represent the mean ± s.d., and each data point represents an individual rat. For (M-N) and (P-U), *P* values were calculated by two-tailed Student’s t-tests. See also Figure S7.

To generate a rat model of Leigh syndrome, we tested eTd-mtABEs targeting T_7_ of mitochondrial *ND6* site 4 to induce m.13875 (corresponding to human m.14487) T-to-C mutation in PC12 cells (Figure 7H). In R-AD version with minimal bystander mutation, compared to sTALED which induced ∼20% editing, all eTd6-mtABEs achieved high mutation rates of 53-69% with nearly single-base T_7_-to-C editing. Then, the eTd6-mtABE-RW/V28A construct was selected for rat zygotes injection (Figures 7I and S7E). HTS data revealed robust desired editing ranging from 26-74% (54% on average) and minimal bystander mutations in all 49 F0 rats (Figures 7J-7L and S7A). The m.13875 T-to-C mutation was efficiently transmitted to the F1 generation (Figure S7F). Analysis of the editing outcomes at different organs from founders showed similar but variable mitochondrial adenine conversion ratios (Figure S7G).

As Leigh syndrome in human is associated in skeletal muscle and heart defects,^43,47^ we leveraged the Rotarod Test and Grip Strength Test to assess the founder rats. The results indicated dramatical impairments in motor coordination, body balance and forelimb grip strength in the founders (Figures 7M, 7N, S7H and S7I). Furthermore, echocardiography confirmed abnormal cardiac structure and function, including dilated heart chambers, increased left ventricular end-systolic volume (LVVS), and decreased left ventricular ejection fraction (LVEF) (Figures 7O-7U and S7J). Intriguingly, we also observed that gender-biased phenotypic manifestations of the disorder, female founders were more susceptible to muscle defects, whereas cardiac abnormalities were more prominent in male founders (Figures S7H-S7J). These data demonstrate that eTd-mtABEs can induce robust *in vivo* mtDNA editing to successfully generate mitochondrial disease-related rat models.

## Discussion

Mitochondrial adenine base editors are crucial tools for mtDNA disease modeling and potentially for gene therapy, as A•T-to-G•C editing can theoretically model and correct over 40% of mtDNA diseases.^3,21^ However, it requires a very high ratio of base conversion frequencies (typically >50%).^5,25^ In this study, we have evolved hyperactive TadA variants to increase the activity of both nuclear and mitochondrial base editors, especially in non-canonical editing windows and broad sequence contexts.

As TadA prefer YA* sequence context especially outside the major editing window,^26,30,31^ we believed that this could be an important start point to evolve TadA-8e mutants to reach higher activity. We demonstrated that A142W or A142R/L145W mutation can enhance the activity and expand the editing window regardless of sequence context in both nuclear and mitochondrial ABEs. Our previous study showed that L145 was a critical residue for substrate recognition, as L145T/Q/C substitution narrowed the editing window, reduced off-target effects and cytosine bystander edits.^33^ Here we found that L145W increased activity but reduced cytosine editing effects (Figures 2B and S2C). Comparing to T/Q/C, the bulky mutation of L145W may pack against F84 and adjust L145 to an even more optimal position for deamination. An additional mutation at the nearby position (A142R) may further strengthen the hydrophobic patch created by W145 and F84. Without the L145W mutation, the single A142W mutation may function similarly to L145W. Although TadA7.10 is screened from TA preference targets,^26^ we have demonstrated that extensive evolution can push the deaminase evolving to more broad targeting scope. During the course of our study, an independent group reported the development of TadA8r which was evolved from wild-type *E.coli* TadA to increase context compatibility in nuclear DNA.^48^ The authors considered that the different substitution at D108 residue (D108G in TadA8r and D108N in TadA7.10/8e descendants) led their base editors to evolve along a distinct trajectory. Our study shows that the D108G substitution is not necessarily required to relax TadA sequence preference, and will contribute to the understanding of the working mechanisms of deaminase and further evolution of base editors.

Compared to their original versions, higher adenine conversion activity was observed in mitochondarial ABEs (up to average 6.4-fold in RA* context and 2.4-fold in YA* context) in comparation to nuclear ABEs (up to average 4.0-fold in RA* context and 1.9-fold in YA* context) (Figures 2D and 3K), suggesting that enhancement of deaminase activity is more effective in mitochondrial ABEs. This is probably due to the different mechanisms for exposing the ssDNA substrate by Cas9 and DddA, as Cas9 functions as a helicase and exposes the non-target strand more thoroughly assisted by sgRNA.^49,50^ It is evidenced that the evolved DddA6 variant,^39^ which showed higher activity and an expanded targeting scope in DdCBEs, could further increase base editing efficiency at most tested sites but not all (Figures 3E, 3G, 6B and S4B). Given the more complicated construction of mitochondrial BEs compared to nuclear BEs, it would be important to reveal the underlying rules for generating highly active constructs for specific targets.

Very recently, a V28R variant was identified to reduce RNA off-targets of sTALEDs, but with relatively low on-target activity.^24^ In this study, the additional introduction of either V28A or V106W mutation to eTd-mtABE-RW displayed a significant decrease in off-target mutations at both the DNA and RNA levels. Surprisingly, the accurate version showed increased activity at the tested sites (Figure 4B), since enhanced precision usually led to compromised efficiency.^33,51^ More importantly, besides eTd-mtABE-RW, eTd-mtABE-RW/V28A also enabled highly efficient installation of pathogenic mutations in rat embryos to generate mtDNA disease models, which were valuable resources for both pathological and therapeutic studies. For example, we achieved a very high level of point mutation rate (54% on average) to model Leigh syndrome, and obvious phenotypes were observed in the F0 generation with sexual biased defects. It would be quite interesting to confirm this phenomenon in patients and to further develop gene therapies based on this model, as mitochondrial C-to-T base editors are available.

The development of eTd-mtABEs substantially advances the utility of mitochondrial base editing for disease modeling and potential therapeutic strategies aimed to correct pathogenic point mutations, rather than eliminating mutant mtDNA, which would be dangerous in tissues with extremely high levels of mtDNA mutation.

### Limitations of the study

We have generated hyperactive TadA variants that exhibit enhanced editing activity, regardless of sequence context and greatly reduced DNA and RNA off-target effects for both nuclear and mitochondrial adenine base editors. However, eTd-mtABEs also exhibit considerable bystander editing activity in both mtDNA strands. When using DNA nickase instead of DddA, eTd-mtABEs enhanced strand-biased editing with higher activity, but accuracy and efficiency still need to be improved to achieve single base mtDNA editing. Identifying and utilizing other enzymes to unwind mtDNA strand with a small bulge might be a potential approach to further strengthen precise base conversion in mitochondrial genome.

## Acknowledgments

We thank Y. Zhang from the Flow Cytometry Core Facility of School of Life Sciences in East China Normal University and support from ECNU Public Platform for innovation (011). We thank L. Ji (MedSci) for designing schematic diagrams. This work was partially supported by grants from National Key R&D Program of China (2023YFC3403400 and 2023YFE0209200 to D.L), the National Natural Science Foundation of China (No.32025023, No.32230064 and No.32311530111 to D.L., No.31930016 to W.W., No.31971366 to L.W., No.82230002 to M.L.), grants from the Shanghai Municipal Commission for Science and Technology (21JC1402200 to D.L.), Young Elite Scientist Sponsorship Program by China Association for Science and Technology (2023QNRC001 to L.C.), the Fellowship of China Postdoctoral Science Foundation (No. 8206400139 to Z.Y.) and the support from Lingang Laboratory.

## Author contributions

L.C. and D.L. designed the experiments. L.C., M.H., C.L., M.Y., Y.W., X.G., Y.F., H.H., X.D., H.G., X.C. and L.G. performed the experiments. L.C., M.H., C.L., M.Y., X.G., Y.F., X.D., D.Z., D.M., M.H., Z.Y., M.L, G.S., X.Z., W.W. and D.L. analyzed the data, L.C. and D.L. wrote the manuscript with the input from all the authors. L.C. and D.L. supervised the research.

## Resource availability

### Lead contact

Please direct requests for resources and reagents to Lead Contact, Dali Li(.)

### Materials availability

All unique reagents in this study are available from the Lead Contact and will be provided upon request.

### Data and code availability

Amplicon sequencing data generated during this study are available at the NCBI Sequence Read Archive database under SUB14426488. All data supporting the results are provided in the main text or supplemental information.

Any additional information required to reanalyze the data reported in this work paper is available from the Lead Contact upon request.

## STAR+METHODS

### KEY RESOURCES TABLE

| REAGENT or RESOURCE | SOURCE | IDENTIFIER |
| --- | --- | --- |
| **Bacterial and virus strains** |  |  |
| Trans5α Chemically Competent Cell | TransGen Biotech | Cat#CD201-01 |
| **Chemicals, peptides, and recombinant proteins** |  |  |
| polyethyleneimine | Polysciences | Cat#23966 |
| Lipofectamine LTX | Thermo Fisher Scientific | Cat#15338100 |
| BbsI | Thermo Fisher Scientific | Cat#FD1014 |
| BamHI | Thermo Fisher Scientific | Cat#FD0054 |
| PmeI | Thermo Fisher Scientific | Cat#FD1344 |
| HindIII | Thermo Fisher Scientific | Cat#FD0504 |
| EcoRI | Thermo Fisher Scientific | Cat#FD0275 |
| NotI | Thermo Fisher Scientific | Cat#FD0593 |
| BglII | Thermo Fisher Scientific | Cat#FD0083 |
| NEBridge® Golden Gate Assembly Kit<br>(BsmBI-v2) | New England BioLabs | Cat#E1602 |
| PrimeSTAR® GXL DNA Polymerase | TAKARA | Cat#R051A |
| PrimeSTAR® Max DNA Polymerase | TAKARA | Cat#R045A |
| KOD-Plus-Neo polymerase | TOYOBO | Cat#KOD-401 |
| T4 DNA Ligase | Beyotime Biotech | Cat#D7008 |
| Ampicillin | Sangon Biotech | Cat#A610028 |
| Kanamycin | Sangon Biotech | Cat#A600286 |
| DMEM | Gibco | Cat#10567014 |
| Fetal bovine serum (FBS) | Gibco | Cat#10099141 |
| penicillin and streptomycin (P/S) | Gibco | Cat#15140122 |
| Trypsin | Gibco | Cat#8121407 |
| Agar A | Sangon Biotech | Cat#A600010 |
| NaCl | Sigma | Cat#S5886 |
| Tryptone | Thermo Fisher Scientific | Cat#LP0042 |
| Yeast Extract | Thermo Fisher Scientific | Cat#LP0021 |
| Agarose | Tsingke Biotech | Cat#TSJ001 |
| TRIzol Reagent | Magen Biotech | Cat#R4802 |
| RNase-free Water | TAKARA | Cat#9012 |
| 5M NH4OAc | Thermo Fisher Scientific | Cat#AM9070G |
| M2 medium | Millipore | Cat#MR-015 |
| Dulbecco's Phosphate-Buffered Saline (D-PBS) | Gibco | Cat#1490144 |
| Hyaluronidase | Sigma | Cat#H4272 |
| EmbryoMax® KSOM Mouse Embryo Media | Millipore | Cat#MR-020P-5F |
| Tris | Sangon Biotech | Cat#A600194 |
| 0.5M EDTA, pH 8.0 | Sangon Biotech | Cat#B540625 |
| Acetic Acid Glacial | Titan Biotech | Cat#64-19-7 |
| 20× PBS Buffer | Sangon Biotech | Cat#B548117 |
| Chloroform | SINOPHAIM | Cat#10006818 |
| Dehydrated Alcohol | Titan Biotech | Cat#01101143 |
| isopropyl | Titan Biotech | Cat#01104420 |
| **Critical commercial assays** |  |  |
| Tiagen plasmid mini-extraction kit | Tiagen Biotech | Cat#DP103-03 |
| HiPure Gel Pure Micro Kit | Magen Biotech | Cat#D2110-02 |
| TIANamp Genomic DNA Kit | Tiangen Biotech | Cat# DP304-03 |
| Mitochondria Isolation Kit for Cultured Cells | Thermo Fisher Scientific | Cat#89874 |
| Clone Express II One Step Cloning Kit | Vazyme | Cat#C111-01 |
| Clone Express II Multis One Step Cloning Kit | Vazyme | Cat#C113-02 |
| QuickExtract™ DNA Extraction Solution | Lucigen | Cat#QE0905T |
| Enhanced ATP Assay Kit | Beyotime Biotech | Cat#S0027 |
| ROS Assay Kit | Beyotime Biotech | Cat#S0033S |
| Hifair® II 1st Strand cDNA Synthesis SuperMix | Yeasten | Cat#11120ES60 |
| Pierce™ BCA Protein Assay Kits | Thermo | Cat# A65453 |
| Hi-yield T7 in vitro transcription reagent (N1-Me-pUTP) | HZYMES | Cat#HBP001506 |
| **Deposited data** |  |  |
| Targeted amplicon sequencing data | This study | SUB14426488 |
| **Experimental models: Cell lines** |  |  |
| Human: HEK293T | ATCC | Cat#CRL-3216 |
| Rat: PC12 | ATCC | Cat#CRL-1721 |
| **Experimental model: Organisms/strains** |  |  |
| Sprague-Dawley | Textile | SD |
| **Oligonucleotides** |  |  |
| Primers used for deep sequencing preparation, see Table S1 | This manuscript | N/A |
| **Recombinant DNA** |  |  |
| TALED_Left-ND1-1397C-AD | Addgene | Cat#183892 |
| TALED_Right-ND1-1397N | Addgene | Cat#183898 |
| ND4-DdCBE-left side TALE | Addgene | Cat#157844 |
| ND4-DdCBE-right side TALE | Addgene | Cat#157843 |
| ND4.2-Left TALE-G1397-N-DddA11-mCherry | Addgene | Cat#179682 |
| ND4.2-Right DdCBE-G1397-C-T1413I-GFP | Addgene | Cat#179686 |
| ABE8e | Addgene | Cat#138489 |
| **Software and algorithms** |  |  |
| GraphPad Prism 9.3.1 | GraphPad | https://www.graphpad.com/ |
| Base editing frequencies and indel calculation algorithms | This study | http://www.rgenome.net/be-analyzer/#! |

### EXPERIMENTAL MODEL AND SUBJECT DETAIL

#### Rats

Sprague-Dawley strain rats used in this study were purchased from Shanghai Jihui Textile Technology Co., Ltd. Rats were maintained in specific pathogen-free facilities at 20-22 °C with 40-60% humidity under a 12 h light-dark cycle. All animal experiments met the regulations drafted by the Association for Assessment and Accreditation of Laboratory Animal Care in Shanghai and were approved by the Experimental Animal Welfare Ethics Committee of East China Normal University.

#### Cell lines

Human HEK293T cells (ATCC, CRL-3216) and rat PC12 cells (ATCC, CRL-1721) were cultured in Dulbecco’s modified Eagle’s medium (DMEM, Gibco) supplemented with 10% (v/v) fetal bovine serum (FBS, Gibco) and 1% (v/v) penicillin and streptomycin. All cell types were maintained at 37 °C with 5% CO_2_. HEK293T cells were passaged every 2 or 3 days and PC12 cells were passaged every 3 or 4 days.

### METHOD DETAILS

#### Plasmid construction

The primers used in this study are listed in Table S1. Oligonucleotides are listed in Table S2. Amino acid and nucleotide sequences are listed in Table S3. TALED_Left-ND1-1397C-AD (#183892), TALED_Right-ND1-1397N (#183898), ND4-DdCBE-left side TALE (#157844), ND4-DdCBE-right side TALE (#157843), ND4.2-Left TALE-G1397-N-DddA11-mCherry (#179682), ND4.2-Right DdCBE-G1397-C-T1413I-GFP (#179686) and ABE8e (#138489) plasmids were purchased from Addgene. New nuclear and mitochondrial base editor plasmids were constructed using previously published methods.^52,53^ In brief, DNA fragments were amplified using PrimeSTAR® Max DNA Polymerase (TAKARA) and assembled with a ClonExpress MultiS One Step Cloning Kit (Vazyme) according to the manufacturer’s protocol. For nuclear base editing, TadA-8e variants introduced individual or combinational mutations were constructed by site-directed mutagenesis using a PCR-based method. sgRNAs were constructed by annealing from 95 °C down to room temperature and ligating into BbsI-linearized U6-sgRNA(sp)-EF1α-GFP (Thermo Fisher Scientific). For double-strand mtDNA editing, we firstly construct the original eTd-mtABE expression plasmids (the TALE array was replaced with two inverted BsmBI restriction sites) and assembled the TALE array using Goldengate (New England BioLabs). All TALE array sequences are listed in Table S2. For stand-preferred mtDNA editing, Human codon-optimized MutH and Nt.BspD6I(C) were synthesized (Genewiz). The mitoABE expression plasmids were constructed by fusing MutH/Nt.BspD6I(C) or Tad-8e variants to the C-terminal of TALE array.^22^ For flow cytometry assay, double-strand or stand-preferred mtDNA editing constructs are modified with mCherry or GFP using a P2A sequence. Plasmids were transformed into DH5α chemically competent cells (TransGen Biotech). Plasmids used for transfection were isolated using the Tiangen plasmid mini-extraction kit according to the manufacturer’s instructions.

#### Cell culture and transfection

Human HEK293T cells (ATCC, CRL-3216) and rat PC12 cells (ATCC, CRL-1721) were cultured in Dulbecco’s modified Eagle’s medium (DMEM, Gibco) supplemented with 10% (v/v) fetal bovine serum (FBS, Gibco) and 1% (v/v) penicillin and streptomycin. Both cell types were maintained at 37 °C with 5% CO_2_. HEK293T cells were passaged every 2 or 3 days and PC12 cells were passaged every 3 or 4 days. For nuclear base editing, HEK293T cells were seeded in 24-well plates (Corning) and transfected with 750 ng of nuclear base editor plasmids and 250 ng of sgRNA plasmids at 70-80% confluence using polyethyleneimine (PEI; Polysciences). For evaluating the mitochondrial base editing, HEK293T cells were seeded in 12-well plates (Corning) and transfected with 800 ng of each eTd-mtABE, sTALED, or mitoABE monomer at 70-80% confluence using polyethyleneimine (PEI; Polysciences). PC12 cells were seeded in 12-well plates (Corning) and transfected with 1,600 ng of each eTd-mtABE or TALED monomer at 70-80% confluence using Lipofectamine LTX (Thermo Fisher Scientific).

#### Genomic DNA extraction and amplification

To assess nuclear base editing efficiency, HEK293T cells transfected after 72 h were washed with 1×PBS, trypsinized and collected via centrifugation for genomic DNA extraction. To evaluate mitochondrial base editing efficiency, HEK293T cells transfected after 72 h or PC12 cells transfected after 48 h were washed with 1×PBS and digested with 0.25% trypsin (Gibco) for fluorescence-activated cell sorting (FACS). EGFP- and mCherry-double positive cells were harvested and the genomic DNA was extracted using QuickExtract^TM^ DNA Extraction Solution (Lucigen) according to the manufacturer’s recommended protocol. The extraction solution was incubated at 65 °C for 6 min and then 98 °C for 2 min. To obtain the genotype of modified rat, genomic DNA for PCR was extracted from collected tissues using traditional isopropyl method. Genome loci of interest were amplified with site-specific primers listed in Table S1 using KOD-Plus-Neo DNA Polymerase (TOYOBO).

#### Whole mitochondrial genome sequencing

To evaluate the specificity of eTd-mtABEs in whole mitochondrial genome, HEK293T cells were seeded in 12-well plates (Corning) and transfected with 800 ng of each eTd-mtABE or sTALED monomer at 70-80% confluence using polyethyleneimine. HEK293T cells transfected after 72 h were washed with 1×PBS, trypsinized and collected via centrifugation at 1,000 rpm, 4 °C. The mitochondria were isolated from transfected cells using the Mitochondria Isolation Kit for Cultured Cells (Thermo Fisher Scientific) according to the manufacturer’s protocol. mtDNA was extracted from isolated mitochondria using the TIANamp Genomic DNA Kit. Long-range PCR was amplified by PrimeSTAR® GXL polymerase (TAKARA) using two sets of partially overlapping primers (listed in Table S1) to capture the whole mtDNA genome. PCR products were purified using HiPure Gel Pure Micro Kit (Magen Biotech). Subsequently, the whole genome sequencing was performed at the mean coverage of 13151× by Illumina NovaSeq 6000 platform (Genewiz). Regions of interest were amplified by PCR with site-specific primers listed in Table S1.

#### RNA purification and targeted RNA sequencing

HEK293T cells were seeded in 12-well plates (Corning) and transfected with 800ng of each eTd-mtABE or sTALED monomer using polyethyleneimine. Cells transfected after 72 h were washed with 1×PBS, trypsinized and collected via centrifugation at 1,000 rpm, 4°C. Collected cells were then homogenized in TRIzol Reagent (Magen Biotech). Total RNA was extracted using standard methods and then reverse transcribed into cDNA using Hifair® Ⅱ 1st Strand cDNA Synthesis SuperMix (Yeasen) according to the manufacturer’s protocol. Genome loci of interest were amplified with site-specific primers listed in Table S1.

#### Intracellular ROS assay

Intracellular ROS assay was measured by a ROS Assay Kit (Beyotime Biotech) following the manufacturer’s instructions. The DCFH-DA probe was diluted to a final concentration of 10μM and Rosup served as the positive control. HEK293T cells were seeded in 12-well plates (Corning) and transfected with 800ng of each eTd-mtABE monomer or L-1397N using polyethyleneimine. After 72 h, the medium was removed and the cells were washed three times with PBS. The staining method was directed by the instructions. In brief, HEK293T cells were incubated with diluted DCFH-DA probe for 5 min at 37 °C and detected through flow cytometry using a BD Fortessa flow cytometer (BD Biosciences).

#### ATP assay

ATP assay was measured by an Enhanced ATP Assay Kit (Beyotime Biotech) following the manufacturer’s instructions. HEK293T cells were seeded in 6-well plates (Corning) and transfected with 800ng of each eTd-mtABE or L-1397N using polyethyleneimine. The transfected Cells after 72 h were washed with 1×PBS and digested with 0.25% trypsin (Gibco) for fluorescence-activated cell sorting (FACS). About 1,000,000 EGFP- and mCherry-double positive cells were harvested and immediately lysed in 200 μl lysis buffer on ice. The protein concentration of each treated group was determined using the Pierce™ BCA Protein Assay Kits (Thermo Fisher Scientific). In a 96-well plate, 20 μl of supernatant was added into the wells containing 100 μL ATP detection working reagent. The plate was then incubated at room temperature for 5 min. The luminescence was detected by EnVision Multilabel Reader and the total ATP levels were defined as nmol/mg.

#### mRNA preparation

IVT template DNAs were prepared by linearizing with EcoRI and extracted by phenol chloroform extracting method. mRNAs were synthesized using Hi-yield T7 in vitro transcription reagent (N1-Me-pUTP) (HZYMES) according to the manufacturer’s protocol. In brief, after 2h incubated at 37 °C, 1 ul of DNase I was added into the solution and incubated at 37 °C for 15 minutes to digest the template DNA. Subsequently, IVT reaction solution was purified by ammonium acetate and washed with pre-cooling 70% ethanol. mRNA was eluted in RNase-free Water (TAKARA) and stored at −80 °C.

#### Animals and microinjection of zygotes

Animal manipulations were in line with a previous report.^33^ The mixture of eTd-mtABE-encoding mRNAs (150 ng/mL each) was diluted in RNase-free Water and injected into cytoplasm using an Eppendorf TransferMan NK2 micromanipulator. Injected zygotes were transferred into pseudo-pregnant female rats at 7-8 weeks old.

#### Rotarod test

To assess the motor coordination of rats, a rotarod machine with automatic timers and falling sensor (XR-6D, Shanghai Xinruan Information Technology) was used. Rats were acclimated to the rotarod at a constant speed of 4 rpm/min for 10 minutes to training. Rotarod test was started at an initial speed of 4 rpm/min and accelerated uniformly to 40 rpm/min within 5 minutes. The time rats spent on rotarod and the rotarod speed rats fell was recorded.

#### Grip test

To evaluate the muscle strength of rats, the maximal force of the forelimbs was measured using a BIO-GS3 (Bioseb) according to the manufacturer’s instructions. The rat hung on a metal bar with its forepaws until the grip failed, while the tester pulled the tail of the rat. Three repeats were performed on each rat, and the average data were calculated.

#### Auditory brainstem response (ABR) measurement

The ABR measurements were consistent with a previous report in a shielded, double-walled sound room.^54^ Rats were anesthetized with intraperitoneal injection of pentobarbitone (50 mg/kg of body weight), and three electrodes were inserted into the subcutaneous tissues at the scalp midline (the recording electrode), posterior to the stimulated ear (the reference electrode), and on the midline of the back 1-2 cm posterior to the neck of the animal (the ground electrode). Tone pips (3, 10, 15 and 20 kHz) at different intensities were generated and delivered using TDT System III (Tucker-Davis Technologies) to test the frequency-specific hearing thresholds. Each sound stimulus was played 20 times per second for 10 seconds and passed through the sound guide tube into the rat’s external ear canal. The ABR signals were acquired, filtered, amplified, and analyzed using equipment and software (BioSig) manufactured by Tucker-Davis Technologies. The ABR threshold was defined as the lowest sound intensity capable of eliciting a response pattern characteristic of that observed at higher intensities. The animal’s body temperature was monitored using a rectal probe and maintained at ∼37 °C by a feed-back-controlled heating blanket.

#### Echocardiography analysis

The heart structure and function of mutant rats were evaluated by echocardiography. The mutant rats were in lightly anesthetized (heart rate ≥370 bpm) and compared with same-aged wild-type rats. Echocardiographic observations were performed using Vevo LAZR-X ultrasound machine with an MX250S probe (20 MHz). M-mode images were used to measure end-systole (LVIDS), end-diastole (LVIDD), left ventricular volume at end-systole and end-diastole (LVVS, LVVD), left ventricular percent fractional shortening (LVFS) and left ventricular ejection fraction (LVEF). Measurements were recorded at least three continuous cardiac cycles.

#### Next-generation sequencing (NGS) and data analysis

The second PCR amplifications were performed with primers containing an adaptor sequence (forward 5’-GGAGTGAGTACGGTGTGC-3’; backward 5’-GAGTTGGATGCTGGATGG-3’) and diverse barcode sequences at the 5′ end. The resulting High-throughput sequencing (HTS) libraries were pooled and purified by electrophoresis with a 1.5% agarose gel using HiPure Gel Pure DNA Micro Kit (Magen) eluting with 60 μl H_2_O, and then sequenced on an Illumina HiSeq platform. To assess base editing efficiencies, A•T-to-G•C efficiencies and indels in the HTS data were analyzed using BE-Analyzer.^55^ Base editing efficiencies were calculated as: base substitution reads divided by total reads. Purities were calculated as: percentage of (the reads of A•T-to-G•C edits) / (the reads of adenine edits without indels). Indel frequencies were calculated as: percentage of (the reads of insertions and deletions) / (total reads).

#### Analysis of mitochondrial genome-wide off-target editing

The analysis of whole mitochondrial genome sequencing data was performed as previously reported.^3^ Initially, we aligned the Fastq sequences to the GRCh38 (release v102) reference genome using BWA (v.0.7.17), and then created BAM files with SAMtools (v.1.9) by fixing read pairing information and flags. Subsequently, we utilized the REDItoolDenovo.py script from REDItools (v.1.2.1) to find all thymines and adenines in the mitochondrial genome with conversion rates > 0.1%. Positions with conversion rates ≥10% in both treated and untreated samples were identified as SNVs in the cell lines and removed. We also excluded the construct’s on-target sites. The remaining sites were regarded as off-target sites, and we counted the number of edited A/T nucleotides with editing frequencies > 0.1%. We calculated the average A•T-to-G•C editing frequency for all bases in the mitochondrial genome by averaging the conversion rates at each base location in the off-target sites. Average mtDNA-wide A-to-G editing frequency was calculated as: percentage of (the sum of A•T-to-G•C off-target) / (all bases in the mitochondrial genome). Average mtDNA-wide C-to-T editing frequency was calculated as: percentage of (the sum of C•G-to-T•A off-target) / (all bases in the mitochondrial genome). Mitochondrial genome-wide graphs were constructed by plotting the conversion rates at on-target and off-target sites with an editing frequency ≥1% across the entire mitochondrial genome.

#### Statistical analysis and reproducibility

Data are presented as mean ± s.d. from independent experiments. All statistical analyses were performed on n = 3 biologically independent experiments unless otherwise noted in the figure captions, using GraphPad Prism version 9.3.1 software.

## References

1. Wallace, D.C. (2018). Mitochondrial genetic medicine. Nat. Genet. 50, 1642–1649. 10.1038/s41588-018-0264-z.

2. Kim, J.S., and Chen, J. (2024). Base editing of organellar DNA with programmable deaminases. Nat. Rev. Mol. Cell Biol. 25, 34–45. 10.1038/s41580-023-00663-2.

3. Cho, S.I., Lee, S., Mok, Y.G., Lim, K., Lee, J., Lee, J.M., Chung, E., and Kim, J.S. (2022). Targeted A-to-G base editing in human mitochondrial DNA with programmable deaminases. Cell 185, 1764–1776 e1712. 10.1016/j.cell.2022.03.039.

4. Li, G., Li, X., Zhuang, S., Wang, L., Zhu, Y., Chen, Y., Sun, W., Wu, Z., Zhou, Z., Chen, J., et al. (2022). Gene editing and its applications in biomedicine. Sci China Life Sci 65, 660–700. 10.1007/s11427-021-2057-0.

5. Silva-Pinheiro, P., and Minczuk, M. (2022). The potential of mitochondrial genome engineering. Nat. Rev. Genet. 23, 199–214. 10.1038/s41576-021-00432-x.

6. Stewart, J.B. (2021). Current progress with mammalian models of mitochondrial DNA disease. J. Inherit. Metab. Dis. 44, 325–342. 10.1002/jimd.12324.

7. Russell, O.M., Gorman, G.S., Lightowlers, R.N., and Turnbull, D.M. (2020). Mitochondrial Diseases: Hope for the Future. Cell 181, 168–188. 10.1016/j.cell.2020.02.051.

8. Bayona-Bafaluy, M.P., Blits, B., Battersby, B.J., Shoubridge, E.A., and Moraes, C.T. (2005). Rapid directional shift of mitochondrial DNA heteroplasmy in animal tissues by a mitochondrially targeted restriction endonuclease. Proc. Natl. Acad. Sci. U. S. A. 102, 14392–14397. 10.1073/pnas.0502896102.

9. Gammage, P.A., Viscomi, C., Simard, M.L., Costa, A.S.H., Gaude, E., Powell, C.A., Van Haute, L., McCann, B.J., Rebelo-Guiomar, P., Cerutti, R., et al. (2018). Genome editing in mitochondria corrects a pathogenic mtDNA mutation in vivo. Nat. Med. 24, 1691–1695. 10.1038/s41591-018-0165-9.

10. Bacman, S.R., Kauppila, J.H.K., Pereira, C.V., Nissanka, N., Miranda, M., Pinto, M., Williams, S.L., Larsson, N.G., Stewart, J.B., and Moraes, C.T. (2018). MitoTALEN reduces mutant mtDNA load and restores tRNA(Ala) levels in a mouse model of heteroplasmic mtDNA mutation. Nat. Med. 24, 1696–1700. 10.1038/s41591-018-0166-8.

11. Zekonyte, U., Bacman, S.R., Smith, J., Shoop, W., Pereira, C.V., Tomberlin, G., Stewart, J., Jantz, D., and Moraes, C.T. (2021). Mitochondrial targeted meganuclease as a platform to eliminate mutant mtDNA in vivo. Nat. Commun. 12, 3210. 10.1038/s41467-021-23561-7.

12. Rees, H.A., and Liu, D.R. (2018). Base editing: precision chemistry on the genome and transcriptome of living cells. Nat. Rev. Genet. 19, 770–788. 10.1038/s41576-018-0059-1.

13. Mok, B.Y., de Moraes, M.H., Zeng, J., Bosch, D.E., Kotrys, A.V., Raguram, A., Hsu, F., Radey, M.C., Peterson, S.B., Mootha, V.K., et al. (2020). A bacterial cytidine deaminase toxin enables CRISPR-free mitochondrial base editing. Nature 583, 631–637. 10.1038/s41586-020-2477-4.

14. Huang, J., Lin, Q., Fei, H., He, Z., Xu, H., Li, Y., Qu, K., Han, P., Gao, Q., Li, B., et al. (2023). Discovery of deaminase functions by structure-based protein clustering. Cell 186, 3182–3195 e3114. 10.1016/j.cell.2023.05.041.

15. Mi, L., Shi, M., Li, Y.X., Xie, G., Rao, X., Wu, D., Cheng, A., Niu, M., Xu, F., Yu, Y., et al. (2023). DddA homolog search and engineering expand sequence compatibility of mitochondrial base editing. Nat. Commun. 14, 874. 10.1038/s41467-023-36600-2.

16. Guo, J., Yu, W., Li, M., Chen, H., Liu, J., Xue, X., Lin, J., Huang, S., Shu, W., Huang, X., et al. (2023). A DddA ortholog-based and transactivator-assisted nuclear and mitochondrial cytosine base editors with expanded target compatibility. Mol. Cell 83, 1710–1724 e1717. 10.1016/j.molcel.2023.04.012.

17. Sun, H., Wang, Z., Shen, L., Feng, Y., Han, L., Qian, X., Meng, R., Ji, K., Liang, D., Zhou, F., et al. (2023). Developing mitochondrial base editors with diverse context compatibility and high fidelity via saturated spacer library. Nat. Commun. 14, 6625. 10.1038/s41467-023-42359-3.

18. Lim, K., Cho, S.I., and Kim, J.S. (2022). Nuclear and mitochondrial DNA editing in human cells with zinc finger deaminases. Nat. Commun. 13, 366. 10.1038/s41467-022-27962-0.

19. Willis, J.C.W., Silva-Pinheiro, P., Widdup, L., Minczuk, M., and Liu, D.R. (2022). Compact zinc finger base editors that edit mitochondrial or nuclear DNA in vitro and in vivo. Nat. Commun. 13, 7204. 10.1038/s41467-022-34784-7.

20. Lee, S., Lee, H., Baek, G., and Kim, J.S. (2023). Precision mitochondrial DNA editing with high-fidelity DddA-derived base editors. Nat. Biotechnol. 41, 378–386. 10.1038/s41587-022-01486-w.

21. Phan, H.T.L., Lee, H., and Kim, K. (2023). Trends and prospects in mitochondrial genome editing. Exp. Mol. Med. 55, 871–878. 10.1038/s12276-023-00973-7.

22. Yi, Z., Zhang, X., Tang, W., Yu, Y., Wei, X., Zhang, X., and Wei, W. (2023). Strand-selective base editing of human mitochondrial DNA using mitoBEs. Nat. Biotechnol. 10.1038/s41587-023-01791-y.

23. Hu, J., Sun, Y., Li, B., Liu, Z., Wang, Z., Gao, Q., Guo, M., Liu, G., Zhao, K.T., and Gao, C. (2023). Strand-preferred base editing of organellar and nuclear genomes using CyDENT. Nat. Biotechnol. 10.1038/s41587-023-01910-9.

24. Cho, S.I., Lim, K., Hong, S., Lee, J., Kim, A., Lim, C.J., Ryou, S., Lee, J.M., Mok, Y.G., Chung, E., et al. (2024). Engineering TALE-linked deaminases to facilitate precision adenine base editing in mitochondrial DNA. Cell 187, 95–109 e126. 10.1016/j.cell.2023.11.035.

25. Gorman, G.S., Chinnery, P.F., DiMauro, S., Hirano, M., Koga, Y., McFarland, R., Suomalainen, A., Thorburn, D.R., Zeviani, M., and Turnbull, D.M. (2016). Mitochondrial diseases. Nat Rev Dis Primers 2, 16080. 10.1038/nrdp.2016.80.

26. Gaudelli, N.M., Komor, A.C., Rees, H.A., Packer, M.S., Badran, A.H., Bryson, D.I., and Liu, D.R. (2017). Programmable base editing of A*T to G*C in genomic DNA without DNA cleavage. Nature 551, 464–471. 10.1038/nature24644.

27. Zhou, C., Sun, Y., Yan, R., Liu, Y., Zuo, E., Gu, C., Han, L., Wei, Y., Hu, X., Zeng, R., et al. (2019). Off-target RNA mutation induced by DNA base editing and its elimination by mutagenesis. Nature 571, 275–278. 10.1038/s41586-019-1314-0.

28. Grunewald, J., Zhou, R., Iyer, S., Lareau, C.A., Garcia, S.P., Aryee, M.J., and Joung, J.K. (2019). CRISPR DNA base editors with reduced RNA off-target and self-editing activities. Nat. Biotechnol. 37, 1041–1048. 10.1038/s41587-019-0236-6.

29. Rees, H.A., Wilson, C., Doman, J.L., and Liu, D.R. (2019). Analysis and minimization of cellular RNA editing by DNA adenine base editors. Sci. Adv. 5, eaax5717. 10.1126/sciadv.aax5717.

30. Richter, M.F., Zhao, K.T., Eton, E., Lapinaite, A., Newby, G.A., Thuronyi, B.W., Wilson, C., Koblan, L.W., Zeng, J., Bauer, D.E., et al. (2020). Phage-assisted evolution of an adenine base editor with improved Cas domain compatibility and activity. Nat. Biotechnol. 38, 883–891. 10.1038/s41587-020-0453-z.

31. Gaudelli, N.M., Lam, D.K., Rees, H.A., Sola-Esteves, N.M., Barrera, L.A., Born, D.A., Edwards, A., Gehrke, J.M., Lee, S.J., Liquori, A.J., et al. (2020). Directed evolution of adenine base editors with increased activity and therapeutic application. Nat. Biotechnol. 38, 892–900. 10.1038/s41587-020-0491-6.

32. Chen, L., Zhu, B., Ru, G., Meng, H., Yan, Y., Hong, M., Zhang, D., Luan, C., Zhang, S., Wu, H., et al. (2023). Re-engineering the adenine deaminase TadA-8e for efficient and specific CRISPR-based cytosine base editing. Nat. Biotechnol. 41, 663–672. 10.1038/s41587-022-01532-7.

33. Chen, L., Zhang, S., Xue, N., Hong, M., Zhang, X., Zhang, D., Yang, J., Bai, S., Huang, Y., Meng, H., et al. (2023). Engineering a precise adenine base editor with minimal bystander editing. Nat. Chem. Biol. 19, 101–110. 10.1038/s41589-022-01163-8.

34. Jeong, Y.K., Lee, S., Hwang, G.H., Hong, S.A., Park, S.E., Kim, J.S., Woo, J.S., and Bae, S. (2021). Adenine base editor engineering reduces editing of bystander cytosines. Nat. Biotechnol. 39, 1426–1433. 10.1038/s41587-021-00943-2.

35. Tu, T., Song, Z., Liu, X., Wang, S., He, X., Xi, H., Wang, J., Yan, T., Chen, H., Zhang, Z., et al. (2022). A precise and efficient adenine base editor. Mol. Ther. 30, 2933–2941. 10.1016/j.ymthe.2022.07.010.

36. Lapinaite, A., Knott, G.J., Palumbo, C.M., Lin-Shiao, E., Richter, M.F., Zhao, K.T., Beal, P.A., Liu, D.R., and Doudna, J.A. (2020). DNA capture by a CRISPR-Cas9-guided adenine base editor. Science 369, 566–571. 10.1126/science.abb1390.

37. Arbab, M., Shen, M.W., Mok, B., Wilson, C., Matuszek, Z., Cassa, C.A., and Liu, D.R. (2020). Determinants of Base Editing Outcomes from Target Library Analysis and Machine Learning. Cell 182, 463–480 e430. 10.1016/j.cell.2020.05.037.

38. Kim, H.S., Jeong, Y.K., Hur, J.K., Kim, J.S., and Bae, S. (2019). Adenine base editors catalyze cytosine conversions in human cells. Nat. Biotechnol. 37, 1145–1148. 10.1038/s41587-019-0254-4.

39. Mok, B.Y., Kotrys, A.V., Raguram, A., Huang, T.P., Mootha, V.K., and Liu, D.R. (2022). CRISPR-free base editors with enhanced activity and expanded targeting scope in mitochondrial and nuclear DNA. Nat. Biotechnol. 40, 1378–1387. 10.1038/s41587-022-01256-8.

40. Yan, D., Ren, B., Liu, L., Yan, F., Li, S., Wang, G., Sun, W., Zhou, X., and Zhou, H. (2021). High-efficiency and multiplex adenine base editing in plants using new TadA variants. Mol Plant 14, 722–731. 10.1016/j.molp.2021.02.007.

41. Catarino, C.B., Ahting, U., Gusic, M., Iuso, A., Repp, B., Peters, K., Biskup, S., von Livonius, B., Prokisch, H., and Klopstock, T. (2017). Characterization of a Leber’s hereditary optic neuropathy (LHON) family harboring two primary LHON mutations m.11778G>A and m.14484T>C of the mitochondrial DNA. Mitochondrion 36, 15–20. 10.1016/j.mito.2016.10.002.

42. Macmillan, C., Kirkham, T., Fu, K., Allison, V., Andermann, E., Chitayat, D., Fortier, D., Gans, M., Hare, H., Quercia, N., et al. (1998). Pedigree analysis of French Canadian families with T14484C Leber’s hereditary optic neuropathy. Neurology 50, 417–422. 10.1212/wnl.50.2.417.

43. Thorburn, D.R., Rahman, J., and Rahman, S. (1993). Mitochondrial DNA-Associated Leigh Syndrome and NARP. In GeneReviews((R)), M.P. Adam, J. Feldman, G.M. Mirzaa, R.A. Pagon, S.E. Wallace, L.J.H. Bean, K.W. Gripp, and A. Amemiya, eds.

44. Khoo, A., Naidu, S., Wijayendran, S.B., Merve, A., Bremner, F., and Sidhu, M.K. (2021). Progressive myoclonic epilepsy due to rare mitochondrial ND6 mutation, m.14487T>C. BMJ Neurol Open 3, e000180. 10.1136/bmjno-2021-000180.

45. Kytovuori, L., Gardberg, M., Majamaa, K., and Martikainen, M.H. (2017). The m.7510T>C mutation: Hearing impairment and a complex neurologic phenotype. Brain Behav 7, e00859. 10.1002/brb3.859.

46. Mutai, H., Watabe, T., Kosaki, K., Ogawa, K., and Matsunaga, T. (2017). Mitochondrial mutations in maternally inherited hearing loss. BMC Med. Genet. 18, 32. 10.1186/s12881-017-0389-4.

47. Dermaut, B., Seneca, S., Dom, L., Smets, K., Ceulemans, L., Smet, J., De Paepe, B., Tousseyn, S., Weckhuysen, S., Gewillig, M., et al. (2010). Progressive myoclonic epilepsy as an adult-onset manifestation of Leigh syndrome due to m.14487T>C. J Neurol Neurosurg Psychiatry 81, 90–93. 10.1136/jnnp.2008.157354.

48. Xiao, Y.L., Wu, Y., and Tang, W. (2024). An adenine base editor variant expands context compatibility. Nat. Biotechnol. 10.1038/s41587-023-01994-3.

49. Jiang, F., and Doudna, J.A. (2017). CRISPR-Cas9 Structures and Mechanisms. Annu Rev Biophys 46, 505–529. 10.1146/annurev-biophys-062215-010822.

50. Yin, L., Shi, K., and Aihara, H. (2023). Structural basis of sequence-specific cytosine deamination by double-stranded DNA deaminase toxin DddA. Nat. Struct. Mol. Biol. 30, 1153–1159. 10.1038/s41594-023-01034-3.

51. Kim, Y.B., Komor, A.C., Levy, J.M., Packer, M.S., Zhao, K.T., and Liu, D.R. (2017). Increasing the genome-targeting scope and precision of base editing with engineered Cas9-cytidine deaminase fusions. Nat. Biotechnol. 35, 371–376. 10.1038/nbt.3803.

52. Chen, L., Hong, M., Luan, C., Gao, H., Ru, G., Guo, X., Zhang, D., Zhang, S., Li, C., Wu, J., et al. (2024). Adenine transversion editors enable precise, efficient A*T-to-C*G base editing in mammalian cells and embryos. Nat. Biotechnol. 42, 638–650. 10.1038/s41587-023-01821-9.

53. Zhang, X., Chen, L., Zhu, B., Wang, L., Chen, C., Hong, M., Huang, Y., Li, H., Han, H., Cai, B., et al. (2020). Increasing the efficiency and targeting range of cytidine base editors through fusion of a single-stranded DNA-binding protein domain. Nat. Cell Biol. 22, 740–750. 10.1038/s41556-020-0518-8.

54. Cheng, Y., Tang, B., Zhang, G., An, P., Sun, Y., Gao, M., Zhang, Y., Shan, Y., Zhang, J., Liu, Q., et al. (2022). Degraded cortical temporal processing in the valproic acid-induced rat model of autism. Neuropharmacology 209, 109000. 10.1016/j.neuropharm.2022.109000.

55. Hwang, G.H., Park, J., Lim, K., Kim, S., Yu, J., Yu, E., Kim, S.T., Eils, R., Kim, J.S., and Bae, S. (2018). Web-based design and analysis tools for CRISPR base editing. BMC Bioinformatics 19, 542. 10.1186/s12859-018-2585-4.

